# Population structure analysis and laboratory monitoring of *Shigella* with a standardised core-genome multilocus sequence typing scheme

**DOI:** 10.1101/2021.06.08.447214

**Authors:** Iman Yassine, Sophie Lefèvre, Elisabeth E. Hansen, Corinne Ruckly, Isabelle Carle, Monique Lejay-Collin, Laëtitia Fabre, Rayane Rafei, Dominique Clermont, Maria Pardos de la Gandara, Fouad Dabboussi, Nicholas R. Thomson, François-Xavier Weill

**Author notes:** Harvard Medical School, Boston, 02115, United States (Elisabeth E. Hansen). Corresponding author (F.-X. Weill).

## Abstract

The laboratory surveillance of bacillary dysentery is based on a standardised *Shigella* typing scheme that classifies *Shigella* strains into four serogroups and more than 50 serotypes on the basis of biochemical tests and lipopolysaccharide O-antigen serotyping. Real-time genomic surveillance of *Shigella* infections has been implemented in several countries, but without the use of a standardised typing scheme. We studied over 4,000 reference strains and clinical isolates of *Shigella*, covering all serotypes, with both the current serotyping scheme and the standardised EnteroBase core-genome multilocus sequence typing scheme (cgMLST). The *Shigella* genomes were grouped into eight phylogenetically distinct clusters, within the *E. coli* species. The cgMLST hierarchical clustering (HC) analysis at different levels of resolution (HC2000 to HC400) recognised the natural groupings for *Shigella*. By contrast, the serotyping scheme was affected by horizontal gene transfer, leading to a conflation of genetically unrelated *Shigella* strains and a separation of genetically related strains. The use of this cgMLST scheme will enhance the laboratory surveillance of *Shigella* infections.

## INTRODUCTION

*Shigella* belongs to the *Enterobacteriaceae* family, and causes bacillary dysentery, a common cause of diarrhoea in low- and middle-income countries. It has been estimated that this intracellular human pathogen, which is transmitted via the faecal-oral route with very low infectious dose (10-100 cells), is responsible for over 210,000 deaths per year, mostly in children under the age of five years^1–3^. In high-income countries, *Shigella* infections also occur in travellers and in some high-risk groups, such as men who have sex with men (MSM) and Orthodox Jewish communities^2–5^. The morbidity of these infections is currently increasing due to growing resistance to antimicrobial drugs in these bacteria^2,3,5,6^.

Laboratory surveillance of *Shigella* infections was initiated several decades ago, and was facilitated by the adoption of a standardised *Shigella* typing scheme in the late 1940s^7^. This scheme, which is still in use today, is based on biochemical tests and serotyping (slide agglutination with typing sera directed against the different *Shigella* lipopolysaccharide O-antigens). It splits the *Shigella* genus into four serogroups (originally considered to be species): *Shigella dysenteriae, S. boydii, S. flexneri*, and *S. sonnei*; these four serogroups are then subdivided into more than 50 serotypes. However, modern population genetics methods, such as multilocus sequence typing (MLST) analysis, and, more recently, core-genome single-nucleotide variant (cgSNV) analysis, have shown that *Shigella* forms distinct lineages within the species *E. coli*, from which it emerged following the acquisition of a large virulence plasmid (VP) enabling the bacterium to invade intestinal cells^8–11^. In parallel, these host-restricted pathogens converged independently on the *Shigella* phenotype (non-motility, no decarboxylation of lysine, no use of citrate and malonate, and other characteristics, as reported by Pupo and coworkers^8^) through genome degradation. Furthermore, these recent methods have shown that the current typing scheme does not capture the natural groupings of this pathogen^8^. Some molecular data have been taken into account in an update of the *Shigella* serotyping scheme. *S. boydii* 13, for example, was withdrawn from the classification, because it was shown to belong to another species, *Escherichia albertii*, and did not contain the VP^12,13^.

In an increasing number of countries, the laboratory surveillance of *Shigella* infections has now passed from conventional serotyping to real-time genomic surveillance^10,14^. The genomic methods used were developed recently, and most of their targets lie within the O-antigen gene cluster (*rfb*) or in the *S. flexneri* serotype-converting prophages, to ensure serotype specificity^14,15^. Several other genes in the accessory genome were recently targeted, resulting in the assignment of *Shigella* serotypes to eight clusters^16^. These methods undoubtedly facilitate backward compatibility between the genomic and serotyping data, but do not fully exploit the unprecedented resolution of genomics. An extension of the MLST method to cover a large number of core-genome genes has been developed. This high-resolution method, core-genome MLST (cgMLST), has been successfully used in the surveillance of many pathogens, including *Listeria monocytogenes*^17^ and *Salmonella enterica*^18^. Furthermore, cgMLST data are easy to interpret with clustering threshold methods, such as the hierarchical clustering (HierCC)^19^ implemented in EnteroBase^18^. However, cgMLST has never been used for the comprehensive description of *Shigella* populations, and the utility of this method for the genomic surveillance of *Shigella* infections has not previously been assessed.

In this study, by analysing over 4,000 genomes from phenotypically characterised *Shigella* strains representative of the global diversity of this pathovar of *E. coli*, we aimed: i) to resolve the population structure of *Shigella* by cgMLST, (ii) to create a dictionary of correspondence between cgMLST HC and serotyping data, and (iii) to update the *Shigella* serotyping scheme by describing new serotypes. We demonstrate that the combination of cgMLST HC with *rfb* gene cluster analysis would enhance the laboratory surveillance of *Shigella* infections, while maintaining backward compatibility with the current serotyping scheme.

## METHODS

### Strains selection and typing

In total, 4,187 *Shigella* reference strains and clinical isolates were studied (Supplementary Data 1). The strains and isolates originated from the French National Reference Centre for *E. coli, Shigella*, and *Salmonella* (FNRC-ESS), Institut Pasteur, Paris, except 11 strains belonging to provisional serotypes and provided by the Public Health Agency of Canada, Winnipeg, Canada, the Centers for Disease Control and Prevention. Atlanta, USA, the Tokyo Metropolitan Research Laboratory of Public Health, Tokyo, Japan, and the International Centre for Diarrhoeal Disease Research, Bangladesh, Dhaka. The collection comprised two datasets. The first dataset – the reference dataset – consisted of 317 *Shigella* reference strains covering all the known serotypes – including provisional serotypes – of the four serogroups (at least one strain per serotype); most of the strains studied were historical strains from various geographic locations and time periods. This first dataset included 44 *S. sonnei* from four different lineages, 16 *S. dysenteriae* type 1 and 98 *S. flexneri* serotypes 1 to 5, X and Y, belonging to seven phylogenetic groups (PGs) published in previous studies^2,5,20,21^. The second dataset – the routine dataset – consisted of 3,870 clinical isolates (of the 3,942 isolates received) sequenced by the FNRC-ESS between 2017 and 2020 in the framework of the French national surveillance programme for *Shigella* infections. All these strains and isolates were thoroughly characterised with a panel of biochemical tests and serotyped by slide agglutination assays according to standard protocols, as previously described^22^. Additional typing sera against KIVI 162 and SH-105 were provided by the International Centre for Diarrhoeal Disease Research, and the Public Health Agency of Canada, respectively.

### DNA extraction and sequencing

The total DNA was extracted with the Wizard Genomic DNA Kit (Promega, Madison, WI, USA), the Maxwell 16-cell DNA purification kit (Promega) or the MagNA Pure DNA isolation kit (Roche Molecular Systems, Indianapolis, IN, USA), in accordance with the manufacturer’s recommendations. The 4,187 strains and isolates were sequenced using different Illumina platforms. FqCleanER version 3.0 (https://gitlab.pasteur.fr/GIPhy/fqCleanER) was used to eliminate adapter sequences^23^, correct sequencing errors^24^, and discard low-quality reads. Assemblies were generated with SPAdes^25^ version 3.15.

### Other studied genomes

With the aim of capturing the broadest possible diversity of *Shigella* populations, we searched the *E. coli/Shigella* database in EnteroBase^18^, and selected 81 additional *Shigella* genomes (reference+ dataset) not originating from the Institut Pasteur (Supplementary Methods section “Other studied genomes”). We included 27 enteroinvasive *E. coli* (EIEC) and 68 *E. coli* strains from the ECOR collection (Supplementary Methods section “Other studied genomes”), to place our *Shigella* genomes in the phylogenetic context of the broader diversity of *E. coli*. We also used the closed PacBio sequences available for all *Shigella* serotypes and described by Kim and coworkers^26^, to study the genetic organisation of the *rfb* gene cluster or various operons described in the “Gene analyses” section. However, these closed genomes were not included in the cgMLST analysis, as they were not edited with short reads and the numerous indels in the homopolymers therefore altered the allelic distances (Supplementary Table 1).

### Characterisation of the O-antigen gene clusters

The *Shigella* O-antigen biosynthetic gene (*rfb*) cluster was analysed after extraction of the region between the housekeeping genes *galF* (encoding UTP-glucose-1-phosphate uridylyltransferase) and *gnd* (encoding 6-phosphogluconate dehydrogenase), which are known to flank the *rfb* cluster^27^. Newly identified *rfb* clusters were annotated based on a previously annotated closely matched *E. coli* cluster in the NCBI BLASTn nucleotide collection (nr/nt) database (100% coverage and at least 99% identity) or with ORFfinder (https://www.ncbi.nlm.nih.gov/orffinder/) when no matching cluster was found in the NCBI BLAST database (https://blast.ncbi.nlm.nih.gov/Blast.cgi). The GenBank accession codes of all the *Shigella rfb* clusters are listed in Supplementary Table 2. We also used three tools for *in silico* serotyping: SeroPred, the serotype prediction tool implemented in EnteroBase, ShigaTyper^14^, and ShigEiFinder^16^. Short-read and SPAdes assemblies were used for ShigaTyper and ShigEiFinder, respectively.

### Phylogenetic analyses

We used the *Escherichia/Shigella* cgMLST scheme (2,513 loci) implemented in EnteroBase to study our genomic datasets. The cgMLST sequence types (cgMLST STs) consist of a combination of up to 2,513 integers, each corresponding to a different allele of a core gene, with some missing data (core gene missing, allele not called). Genetic distances between genomes were calculated from the number of shared cgMLST alleles. These bacterial genomes were also assigned to clusters at multiple levels of resolution, by a hierarchical clustering approach (HierCC)^19^ implemented in the “cgMLST V1 + HierCC” tool. The resulting HierCC clusters (HCs), at 13 different, fixed cgMLST allele distances, ranging from HC0 (no allelic difference allowed) to HC2350 (maximum of 2,350 allelic differences) were then assigned stable cluster group numbers. The cgMLST trees were inferred with the NINJA NJ algorithm, based on the “cgMLST V1 + HierCC” scheme. We visualised the cgMLST data with GrapeTree^28^.

We also performed cgSNV analysis, to assess the phylogenetic relationships of 398 *Shigella* (317 from the reference dataset and 81 from the reference+ dataset) and 95 *E. coli* (68 ECOR and 27 EIEC) strains. An *Escherichia fergusonii* genome (RHB19-C05, GenBank accession no. GCF_013892435.1) was used as an outgroup for the cgSNV analysis. The paired-end reads and simulated paired-end reads were mapped onto the reference genome of *E. coli* K12-MG1655 (GenBank accession no. NC_000913.3) with Snippy version 4.6 (https://github.com/tseemann/snippy) using the default parameters but with a minimum read coverage of 4 and a 75% read concordance at a locus for a variant to be reported. An alignment of 92,688 SNVs was used for phylogenetic inference. A maximum-likelihood (ML) phylogenetic tree was generated with RAxML-NG^29^ version 1.0.1 using the general time-reversible (GTR) model of nucleotide substitution with a gamma model of the between-site heterogeneity rate (GTR + G) and 100 bootstrap iterations. The best-scoring ML tree of the 20 replicates was midpoint-rooted and visualised with interactive tree of life (iTOL)^30^ version 6 (https://itol.embl.de).

A phylogenetic tree of *rfb* sequences was constructed with the sequences from 43 *Shigella* (Supplementary Table 2) and 196 *E. coli* strains from DebRoy and coworkers^27^. The *Shigella rfb* sequences were trimmed to ensure the same start and end points as for the *E. coli rfb* sequences from DebRoy and coworkers^27^. A sequence alignment was generated with MEGA X^31^ version 10.2.1, using ClustalW with default settings. A ML phylogeny was created with RAxML-NG^29^ version 1.0.1, using the GTR+G model and 100 bootstrap replicates. The ML tree with the best score of the 20 replicates was midpoint-rooted and visualised with iTOL^30^ version 6 (https://itol.embl.de).

### Gene analyses

The presence of the *ipaH* gene, a multicopy gene unique to *Shigella* and EIEC^32^, the presence and structure of the mannitol (*mtl*)^33^, raffinose^34^, and tryptophanase (*tna*) operons^35^, and the type of the O-antigen gene cluster (*rfb*) were determined on SPAdes assemblies using the NCBI BLASTn tool (https://blast.ncbi.nlm.nih.gov/Blast.cgi). The target sequences are described in Supplementary Table 3.

## RESULTS

### Global population structure of *Shigella*

We assembled and sequenced a collection of 317 *Shigella* strains chosen on the basis of their representativeness of the known diversity of *Shigella* populations (i.e., covering all serogroups and serotypes, and the different lineages or phylogroups of *S. sonnei* and *S. flexneri*). The genomic diversity of this reference dataset was increased further, by adding another 81 publicly available *Shigella* genomes. The 398 genomes studied were from strains belonging to the *S. flexneri* (*n* = 191), *S. dysenteriae* (*n* = 83), *S. boydii* (*n* = 80), and *S. sonnei* (*n* = 44) serogroups (Supplementary Table 4). We determined the wider phylogenetic context of these *Shigella* genomes, by also analysing 95 *E. coli* genomes, including 27 EIEC from eight different EIEC genomic clusters and 68 (of the 72) strains from the ECOR collection, considered representative of the diversity of natural populations of *E. coli*^36^. These 493 genomes were studied by two different approaches: the EnteroBase *Escherichia/Shigella* cgMLST scheme and SNV-based clustering.

According to cgMLST, all these genomes belonged to the same hierarchical cluster, HC2350_1 (Supplementary Data 1). As expected, all the *Shigella* and EIEC genomes contained the pathogenicity gene *ipaH*, whereas the ECOR genomes did not (Supplementary Fig. 1). A NINJA neighbour-joining (NJ) tree of core genomic allelic distances was generated with the dataset for the 493 *Shigella* and *E. coli* genomes (Fig. 1A). The differential contribution of the reference and reference+ datasets to *Shigella* population diversity is shown in Supplementary Fig. 2. Visual examination of the colour-coded HC2000 tree revealed that the *Shigella* genomes were grouped into eight different HC2000 clusters (Fig. 1B). Seven of these HC2000 clusters contained exclusively *Shigella* genomes. The eighth, HC2000_2, contained *S. dysenteriae* type 8 and *E. coli* (EIEC and ECOR) genomes. Four HC2000 clusters contained *Shigella* genomes from a single serotype: HC2000_305 (*S. sonnei*), HC2000_1463 (*S. dysenteriae* type 1), HC2000_44944 (*S. dysenteriae* 10), and HC2000_45542 (*S. boydii* 12). These clusters are referred to below as SON, SD1, SD10, and SB12, respectively. Three clusters, HC2000_1465, HC2000_4118, and HC2000_192, consisted of multiple serogroups and serotypes (Figs. 1-4). The first of these clusters, HC2000_1465, contained various serotypes of *S. dysenteriae* (3, 4-7, 9, 11-15, provisional (prov.) 93-119, prov. SH-103, prov. 97-10607, prov. SH-105, prov. 96-3162 and prov. 204/96), *S. boydii* (1-4, 6, 8, 10, 11, 14, 18-20, and prov. 07-6597), and *S. flexneri* type 6 (Fig. 2), consistent with Cluster 1 described by Pupo and coworkers^8^ in their MLST analysis of 46 diverse *Shigella* strains. The HC2000_1465 cluster, named S1, can be divided into five HC1100 clusters (Fig. 2). Only the HC1100_36524 cluster (subcluster S1d) contained strains from a single serotype, *S. dysenteriae* 7. The HC1100_45518 cluster (S1e) contained only *S. flexneri* 6 strains, but most strains from this serotype were in another HC1100, HC1100_1465 (S1b), along with *S. dysenteriae* 3 (Supplementary Results section “Aerogenic strains of *S. boydii* 14 and *S. dysenteriae* 3”) and various serotypes of *S. boydii.* The HC1100_1466 cluster (S1c) contained *S. dysenteriae* 5 and various serotypes of *S. boydii*. Finally, the HC1100_4194 cluster (S1a) included only *S. dysenteriae* strains, but from diverse serotypes. *S. dysenteriae* 3 was found in two different S1 subclusters, S1a and S1b. At a higher level of resolution, four *Shigella* serotypes were grouped within specific HC400 clusters, whereas the other serotypes were split between two to six HC400 clusters (Supplementary Table 5).

**Figure 1.**
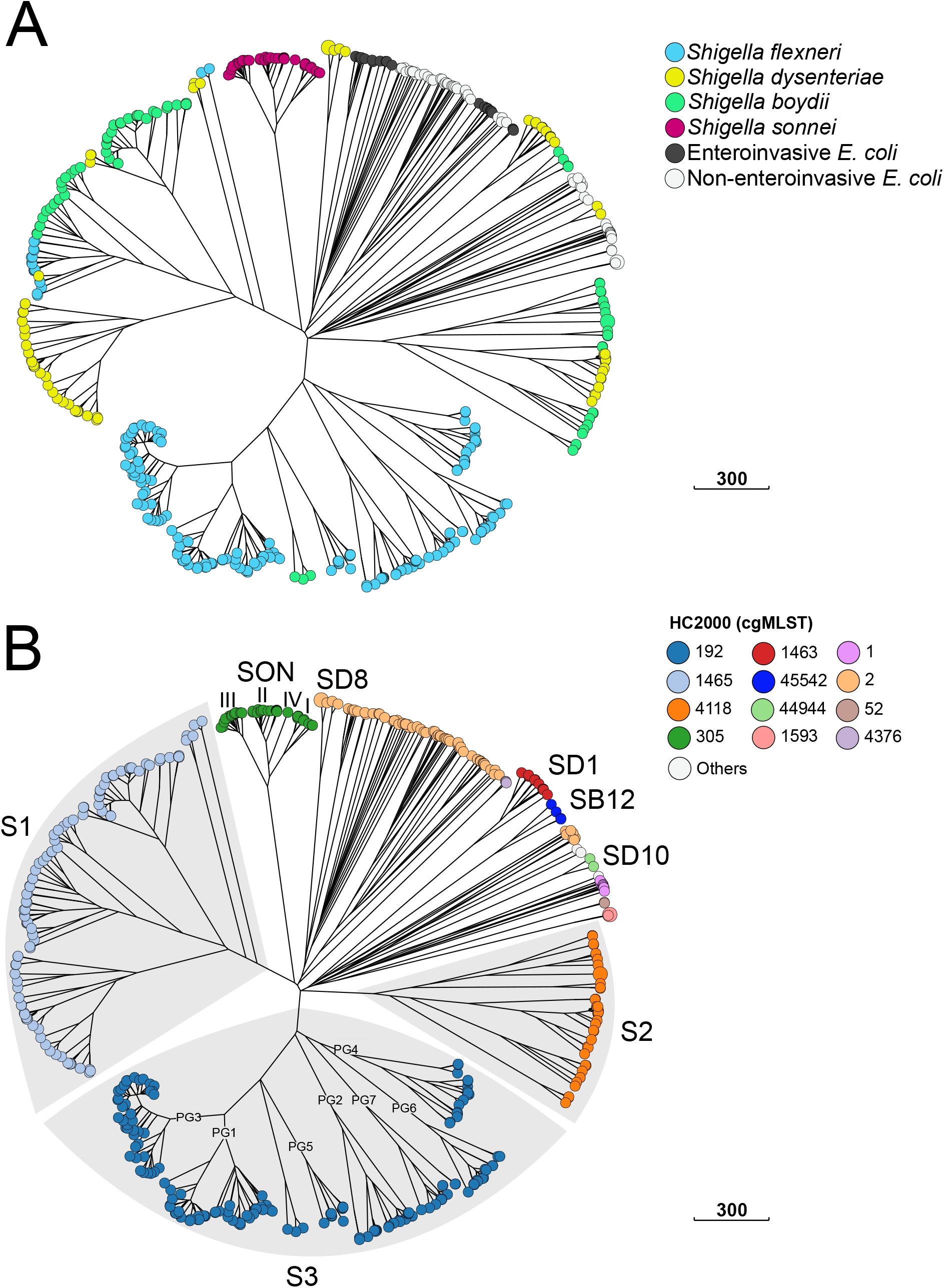
A NINJA neighbour-joining GrapeTree showing the population structure of *Shigella* spp. based on the cgMLST allelic differences between 493 *Shigella* and *E. coli* reference genomes. (A) The tree nodes are colour-coded by *Shigella* serogroup and *E. coli* pathovar. (B) The tree nodes are colour-coded by HC2000 data. HC2000 clusters with fewer than two isolates are represented by white nodes. The different *Shigella* cgMLST clusters are labelled. For the SON cluster, the different genomic lineages of *S. sonnei* are indicated with Latin numerals. For the *S. flexneri* serotypes in cluster S3, the phylogenetic groups (PG1 to PG7) identified by Connor and coworkers^2^ are also indicated. The interactive version of the tree is publicly available from http://enterobase.warwick.ac.uk/ms_tree?tree_id=55118

**Figure 2.**
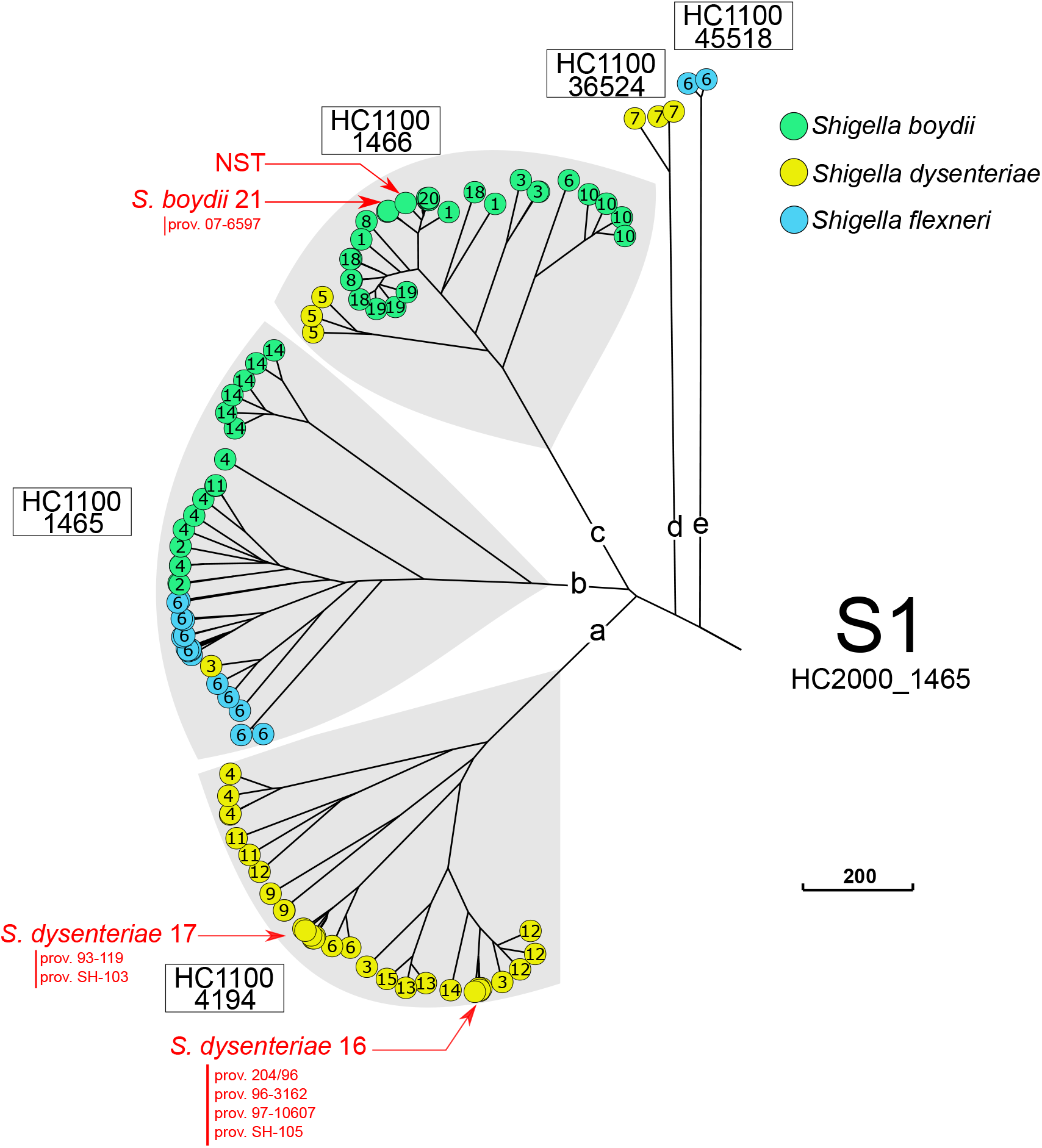
A NINJA neighbour-joining GrapeTree showing the population structure of the *Shigella* S1 cluster (HC2000_1465). This subtree is based on the tree shown in Figure 1. The tree nodes are colour-coded by serogroup. The numbers within nodes indicate the serotype. HC1100 designation is indicated next to each subcluster. Novel and provisional (prov.) *Shigella* serotypes are also shown. NST, = non-serotypable.

The second cluster, HC2000_4118, comprised various serotypes of *S. dysenteriae* (2, prov. E670/74, prov. 96-265, and prov. BEDP 02-5104) and *S. boydii* (5, 7, 9, 11, 15-17) (Fig. 3). This cluster, consisting exclusively of indole-positive strains (Supplementary Results sections “Genomic analysis of the phenotypic markers used in the current *Shigella* typing scheme”), corresponds to the Cluster 2 described by Pupo and coworkers^8^. The HC2000_4118 cluster, hereafter referred to as S2, could be divided into six distinct HC1100 clusters (Fig. 3). Five of these HC1100 clusters contained exclusively *S. boydii*; the sixth, HC1100_4191 (subcluster S2d), contained *S. boydii* 15 and all the *S. dysenteriae* serotypes found in S2. Three HC1100 clusters contained a single serotype: HC1100_11401 (S2f) for *S. boydii* 7, HC1100_7057 (S2e) for *S. boydii* 9, and HC1100_11421 (S2c) for *S. boydii* 11. This last serotype was also found in the S1 cluster (S1b subcluster). At higher resolution, it was possible to assign some serotypes to a particular HC400 cluster. This was the case for *S. boydii* 16 (HC400_11449) and *S. boydii* 17 (HC400_11452). However, at this level of resolution, other serotypes were split between two to four clusters (Supplementary Table 5).

**Figure 3.**
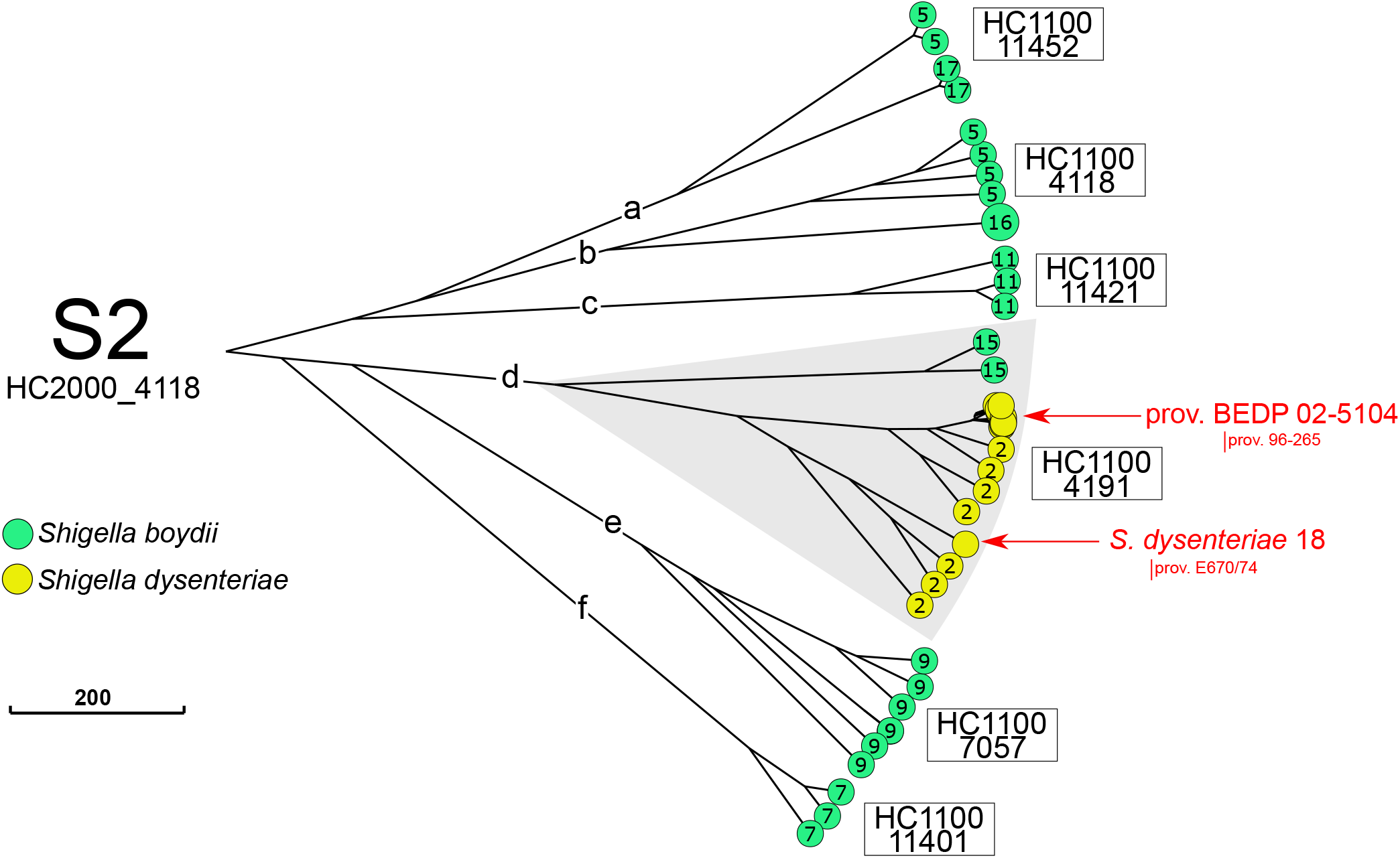
A NINJA neighbour-joining GrapeTree showing the population structure of the *Shigella* S2 cluster (HC2000_4118). This subtree is based on the tree shown in Figure 1. The tree nodes are colour-coded by serogroup. The numbers within nodes indicate the serotype. HC1100 designation is indicated next to each subcluster. Novel and provisional (prov.) *Shigella* serotypes are also shown.

The third cluster, HC2000_192, comprised *S. boydii* prov. E1621-54 (now proposed as *S. boydii* 22; see next section) and all serotypes and subserotypes of *S. flexneri*, with the exception of *S. flexneri* 6, which grouped in S1 (Fig. 4). This cluster seems to correspond to the Cluster 3 reported by Pupo and coworkers^8^, except that *S. boydii* 12 rather than *S. boydii* prov. E1621-54 was reported in Cluster 3 in this previous study (Supplementary Results section “Genomic clustering of *Shigella* reference strains”). This HC2000_192 cluster, hereafter referred to as S3, could be divided into seven distinct HC1100 clusters (Fig. 4A). One of these S3 subclusters, HC1100_11429, contained exclusively *S. boydii* prov. E1621-54. The other six HC1100 clusters contained two or more *S. flexneri* serotypes per cluster. Connor and coworkers^2^ previously subdivided >350 genomes of *S. flexneri* 1-5, X, Y into seven phylogenetic groups (PGs), based on a Bayesian analysis of population structure. As 140 *S. flexneri* genomes from our study were included in the analysis by Connor and coworkers^2^, we compared the clustering by cgMLST HC1100 to that obtained by PG. HC1100_204, HC1100_543, HC1100_1468, HC1100_11594, HC1100_1530 corresponded to PG2, PG4, PG5, PG6 and PG7, respectively (Fig. 4). HC1100_192 encompassed PG1 and PG3, and the use of a higher HC resolution made it possible to link HC400_192 to PG3. However, PG1 did not correspond to a single HC400 cluster. Instead, it corresponded to two such clusters: HC400_237 and HC400_327.

**Figure 4.**
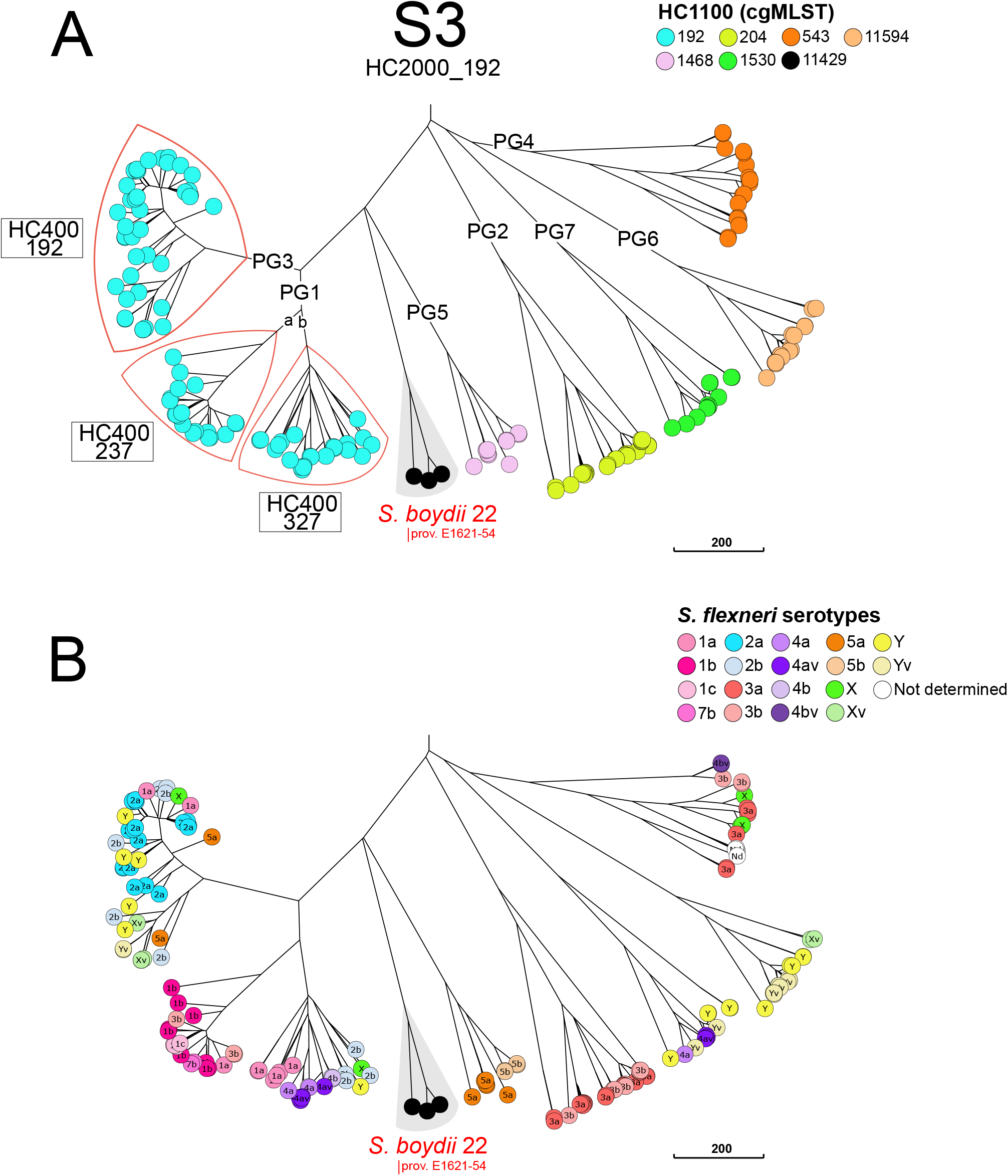
A NINJA neighbour-joining GrapeTree showing the population structure of the *Shigella* S3 cluster (HC2000_192). This subtree is based on the tree shown in Figure 1 (A). The tree nodes are colour-coded by HC1100 data. The *S. flexneri* phylogenetic groups (PG) identified by Connor and coworkers^2^ are indicated. Some HC400 clusters are indicated to separate PG3 from PG1. *S. boydi* 22 (formerly prov. E1621-54) is shown. (B) The tree nodes are colour-coded by *S. flexneri* serotype.

We evaluated the accuracy of cgMLST HC for grouping *Shigella* genomes into different phylogenetic clusters by employing another approach: using the same dataset of 493 *E. coli* and *Shigella* genomes, we constructed a ML tree based on 92,688 SNV differences, and compared this SNV-based clustering (with strong bootstrap support) to the cgMLST HC data. There were no observable differences between the two approaches (Fig. 5).

**Figure 5.**
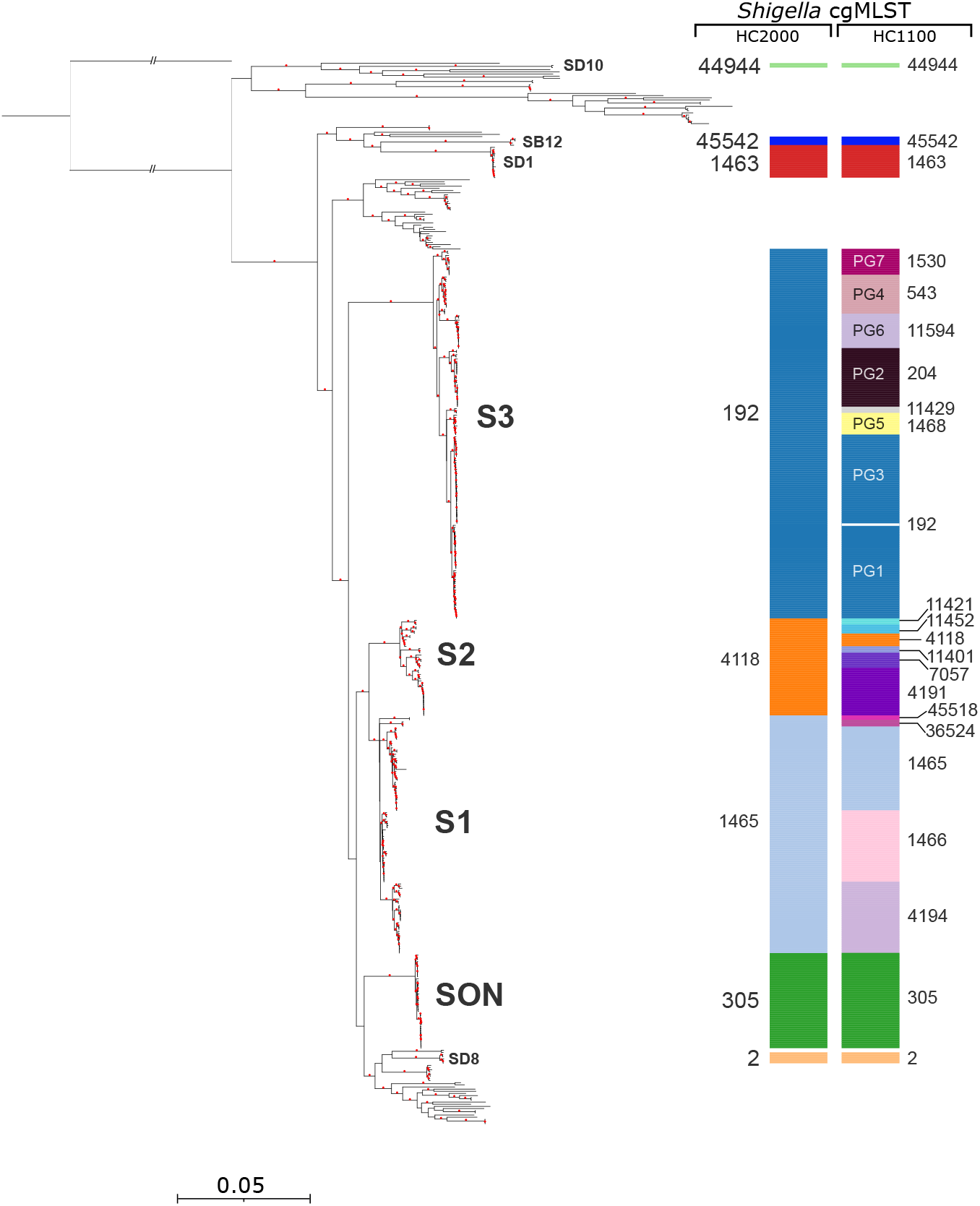
A maximum-likelihood phylogenetic tree showing the population structure of 493 *Shigella* and *E. coli* reference genomes based on 92,688 core-genome single-nucleotide variants (SNVs). Nodes supported by bootstrap values ≥95% are indicated by red dots. Phylogenetic clades containing *Shigella* genomes are labelled with the same nomenclature as in Figure 1. All the *Shigella* genomes are also labelled on the right with cgMLST HC2000 and HC1100 data.

To confirm the robustness of the population structure of *Shigella* established by cgMLST analysis of our reference datasets, we also applied cgMLST to 3,870 clinical *Shigella* isolates received by the FNRC-ESS between 2017 and 2020, in the framework of the French national surveillance programme for *Shigella* infections. All these isolates were characterised phenotypically, on the basis of biochemical reactions and serotyping. They belonged to *S. dysenteriae* (*n* = 53), *S. boydii* (*n* = 101), *S. flexneri* (*n* = 1,555), and *S. sonnei* (*n* = 2,161). All but one of these 3,870 genomes were assigned to known serotype/HC2000/HC1100/HC400 combinations, without inconsistencies (Supplementary Data 1, Supplementary Fig. 2). The exception was an HC1100_204 (PG2) *S. flexneri* isolate, grouped into a new HC400 cluster, HC400_11853.

### Updating the *Shigella* typing scheme

In recent decades, several provisional new serotypes of *S. dysenteriae* and *S. boydii* have been described by different groups across the world^37,38^. However, the phylogenetic relationships between these provisional serotypes and between these serotypes and other *Shigella* populations have not been investigated. We characterised these relationships in detail (Supplementary Results section “Updating the *Shigella* typing scheme”). We found that all these provisional serotypes belonged to the three main *Shigella* clusters, S1 to S3 (Figs. 2-4), and that many of those reported under different names were actually identical. We propose adding *S. dysenteriae* 16-18, and *S. boydii* 21 and 22 to the current serotyping scheme, retaining provisional status for *S. dysenteriae* prov. BEDP 02-5104. All the reference strains for these new serotypes are now available from the *Collection de l’ Institut Pasteur* (CIP) or the National Collection of Type Cultures (NCTC) (Supplementary Results section “Updating the *Shigella* typing scheme”).

### Performance of available *in silico* serotype prediction tools

*In silico* serotyping tools have been developed by various groups, and can be used to maintain links with the current *Shigella* serotyping system. We assessed the performances of the three tools currently available: the EnteroBase “SeroPred” tool^18^, ShigaTyper^14^, and ShigEiFinder^16^ with our 317 genomes from well-characterised reference strains. ShigEiFinder (Supplementary Table 6) gave the best serotype prediction results. However, 100% of the strains belonging to *S. boydii* 10 and to the new serotype *S. dysenteriae* 17, and 14-20% of the strains from *S. boydii* 11, *S. boydii* 14, and *S. dysenteriae* 2 were not identified. All the strains from *S. dysenteriae* prov. BEDP 02-5104 were incorrectly predicted to be *S. dysenteriae* 2, whereas 83% of the strains from the new serotype *S. dysenteriae* 16 were incorrectly predicted to be *S. dysenteriae* prov. 96-265 and 13% were not assigned.

## DISCUSSION

We present here a broad overview of the population of *Shigella*. The hierarchical clustering of cgMLST data and a cgSNV analysis showed that *Shigella* strains belong to eight phylogenetically distinct clusters, within the species *E. coli*. Our results are consistent with previous studies suggesting multiple origins of the *Shigella* phenotype^8,39^. However, the higher resolution of cgMLST, and comprehensive sampling from thousands of phenotypically characterised isolates and reference strains covering all serotypes, including provisional serotypes and atypical strains, made it possible to complete, and in some cases amend, the *Shigella* population structure obtained in previous studies.

The 70-year-old *Shigella* typing scheme, which is still in use today, was based on biochemical characteristics, antigenic relationships, and tradition^7^. We show here that, unlike cgMLST, this scheme does not always reveal natural groupings. In particular, the *Shigella* serogroups/species are artificial constructs developed from data for antigen and metabolic markers affected by Insertion Sequence (IS) element mobilisation and horizontal gene transfer. The presence of large numbers of ISs and their expansions in *Shigella* genomes may alter the nature of both the O-antigen and the rare phenotypic markers identified in this bacterium with weak metabolic activity, by disrupting coding sequences or causing genome rearrangements and deletions^9^. For example, *S. boydii* 6 and 20 arose in subcluster 1c following the acquisition of a single IS within the *rfb* cluster of *S. boydii* 10 and 1, respectively. Serotype diversification, which is observed mostly in clusters S1 to S3, also occurs via horizontal gene transfer of the O antigen-encoding *rfb* cluster from *Escherichia* donors^8,27^. Horizontal gene transfer outside of the *rfb* cluster can also alter the serotype of a strain, as illustrated particularly clearly by the S3 cluster. All the *S. flexneri* strains in this cluster share the same O-antigen backbone structure and their serotypes are determined by glucosylation and/or O-acetylation modifications to the O-antigen tetrasaccharide repeat, conferred by prophage-encoded *gtr* and/or *oac* genes, respectively^15^. Plasmid-mediated serotype conversion by the O-antigen phosphoethanolamine transferase gene (*opt*) has also been reported in *S. flexneri*^15^. Each of the seven *S. flexneri* phylogenetic groups (PGs) described by Connor and coworkers^2^, based on a cgSNV analysis, contained two or more of these serotypes. As this serotyping method does not reflect the genetic relatedness between *Shigella* isolates, and has a number of other disadvantages, including being time-consuming, with intra- and interspecies cross-reactivity, and the impossibility of typing rough strains and new serotypes^14,37^, modern laboratory surveillance of *Shigella* infections should now be based on phylogenetically relevant methods rather than simply on molecular or *in silico* serotyping^10,14-16^. In our hands, the cgMLST HC analysis proved to be the method of choice for monitoring the trends in *Shigella* types. The different types of *Shigella* can be identified with HC2000. Higher resolution, with HC1100 and, in certain cases, HC400, can reveal additional subclusters. This is particularly interesting for S3, which contains the *S. flexneri* 1-5, X, and Y serotypes generated via horizontal gene transfer rather than by vertical descent. We therefore recommend integrating the seven phylogenetic groups (PG1-PG7) described for *S. flexneri* into routine genomic surveillance for *S. flexneri*. These PGs can be easily inferred from cgMLST HC1000/HC400; it is even possible to obtain up to eight groups (after subdividing PG1 into PG1a and PG1b). The cgMLST HC analysis also provides, in a single step, a wide range of clustering levels, from HC0 (no allelic difference allowed) to HC2350 (maximum of 2,350 allelic differences), with a standard nomenclature. For the most frequent *Shigella* serotypes, such as *S. sonnei* and *S. flexneri* 2a, higher resolution levels, such as HC5 and HC10, can also help to identify a single-source outbreak or an epidemic strain, before confirmation by cgSNV analysis. The use of cgMLST HC data also makes it possible to query EnteroBase, which contains over 160,000 *E. coli/Shigella* genomes, to identify strains with similar HC types. This can facilitate the investigation of unusual types of *Shigella* or outbreaks with an international dimension. HC10 was recently used to investigate the origins of an outbreak of *S. sonnei* infections in Belgium, and made it possible to link this outbreak to South America^40^.

However, the use of cgMLST HC data in surveillance should be paired with *in silico* serotyping, to achieve backward compatibility with the current serotyping scheme. This is a very important point for the maintenance of international surveillance with laboratories that cannot currently afford genomic surveillance and to prevent disjunction with the seven decades of serotyping data accumulated worldwide. For this purpose, we found that ShigEiFinder^16^ had the best performance of the three available tools. However, it requires optimisation for certain serotypes. The complete set of *rfb* sequences provided by our study would be helpful for improving this tool.

In conclusion, by studying >4,000 serotyped reference strains and routine isolates covering the overall diversity of *Shigella*, we were able to demonstrate that cgMLST is a robust and portable genomic method revealing natural groupings for this pathovar of *E. coli.* The cgMLST method has strong added value in the framework of the laboratory monitoring of *Shigella*, as it prevents genetically unrelated strains being conflated, and genetically related strains being separated. However, we strongly recommend combining cgMLST with *in silico* serotyping to maintain backward compatibility with the current *Shigella* serotyping scheme.

## SUPPLEMENTARY INFORMATION

Supplementary Information is linked to the online version of the paper at www.nature.com/nature.

## ACKNOWLEDGMENTS

We thank all corresponding laboratories for sending isolates to the French National Reference Centre for *E. coli, Shigella*, and *Salmonella*. We also thank N. Strockbine, K. Talukder, L. Peterson, and S. Matsushita for sending some *Shigella* provisional serotypes, and the sequencing teams at the Institut Pasteur (P2M-Plateforme de Microbiologie Mutualisée) and the Wellcome Sanger Institute for sequencing the samples.

## FUNDING

*Institut Pasteur; Santé publique France*; Fellowship from Association AZM and Saadeh, and the Lebanese Association for Scientific Research (LASeR) for IY. NRT is funded by the Wellcome Trust (grant 206194). The funders had no role in study design, data collection and analysis, decision to publish, or preparation of the manuscript. For the purpose of Open Access, the authors have applied a CC BY public copyright licence to any Author Accepted Manuscript version arising from this submission.

## AUTHOR CONTRIBUTIONS

FXW conceived and designed the study. IY, EH, LF and FXW did the genomic analyses. IY and FXW contributed to data interpretation and visualisation. CR, IC and MLC conducted the laboratory experiments. SL, CR, IC, MLC, MPG, DC, and FXW contributed to isolate acquisition and data collection. FXW, FD, RR and NRT were responsible for funding acquisition. FXW and IY drafted the article. LF, EH, RR, DC, MPG, SL, FD, and NRT critically reviewed the draft. All authors read and approved the final manuscript. IY and FXW accessed and verified the underlying data.

## DATA AVAILABILITY STATEMENT

Short-read sequence data were submitted to EnteroBase^18^ (https://enterobase.warwick.ac.uk/), and to the European Nucleotide Archive (https://www.ebi.ac.uk/ena/) under study numbers PRJEB44801, PRJEB2846, and PRJEB2128. The GrapeTree of the reference and reference+ datasets is publicly available from EnteroBase (http://enterobase.warwick.ac.uk/ms_tree?tree_id=55118). All the GenBank and ENA accession numbers of the genomes used in this study are listed in Supplementary Data 1.

## AUTHOR INFORMATION

Reprints and permissions information is available at www.nature.com/reprints

The authors have no competing financial interests to declare.

Correspondence and requests for materials should be addressed to F.-X.W.

## Notes

### Competing Interest Statement

The authors have declared no competing interest.

