## Supplementary Information for "Population structure analysis and laboratory monitoring of *Shigella* with a standardised core-genome multilocus sequence typing scheme"

### Supplementary Methods

#### Other studied genomes

EnteroBase[1] was queried on 11 November 2020, to identify genomes with new HC1100/HC400 combinations (i.e., not present in our reference and routine datasets) for a given serotype, determined by *in silico* analysis of the *rfb* cluster (see below). We selected at least one genome for each unique combination and a maximum of five if the genomes concerned appeared to come from epidemiologically unrelated isolates. In addition, if a serotype was represented by only one strain in our collection, we selected another genome from EnteroBase, when possible. This selection resulted in 81 *Shigella* genomes (reference+ dataset). The 27 enteroinvasive *E. coli* (EIEC) included in this study belonged to the various previously described EIEC clusters[2–5]. We discarded four (ECOR 7, 23, 32, 43) of the 72 *E. coli* strains from the ECOR collection[6] from this study due to discrepant results for MLST and/or Clermont typing between the studies of Galardini and coworkers[7], Clermont and coworkers[8], and EnteroBase[1]. These 81 additional *Shigella* genomes, the 68 *E. coli* genomes from the ECOR collection, and the 27 EIEC genomes are listed in Supplementary Data 1.

### Supplementary Results

#### Genomic clustering of *Shigella* reference strains

One *Shigella* reference strain originally present in our collection (UE 95-1589)[9] and previously described as *S. boydii* 7 was actually *Escherichia albertii* (HC2350_1596). It did not contain the invasion plasmid antigen gene *ipaH,* instead bearing the intimin *eae* gene, a pathogenicity gene present in enteropathogenic *E. coli* (EPEC) and enterohaemorrhagic *E. coli* (EHEC). This strain, and the *S. boydii* 13 strains, now reclassified as *E. albertii*[10] (HC2350_1596), were not, therefore, included in this study.

The SB12 cluster was not identified by Pupo and coworkers[11], or by Yang and coworkers[12]. Instead, the single *S. boydii* 12 strain studied in each of these studies was assigned to their cluster 3 (our cluster S3), in place of *S. boydii* prov. E1621-54 (not included in their study). As *S. boydii* prov. E1621-54 is serologically related to *S. boydii* 12 (ref. [13]), it seems likely that their strains were actually *S. boydii* prov. E1621-54 but were mistakenly serotyped as *S. boydii* 12 due to the cross-agglutination between the two serotypes. We also encountered this problem when typing *Shigella* isolates collected from children in Niger[14]. The nine indole-negative isolates obtained, reported to be *S. boydii* 12 at the time, were sequenced and found to belong to HC1100_11429, within cluster S3. Following the use of a typing serum against strain E1621-54, they were reclassified as *S. boydii* prov. E1621-54 (now proposed as *S. boydii* 22; see section "Updating the *Shigella* typing scheme"). Furthermore, Pupo and coworkers[11] did not report *S. boydii* serotype 11 in both clusters S1 and S2, but only in S2. A genomic study of 117 *Shigella* isolates belonging to validated serotypes found seven *Shigella* clusters following a SNV-based ML phylogeny analysis[5]. The SB12 cluster was not found, and their *S. boydii* 12 strain instead clustered with their cluster 5 (S2 in our study). The analysis of this genome (SRR2994194) with cgMLST and *rfb* cluster analysis revealed that it was actually *S. dysenteriae* 2. Their *S. boydii* 9 strain was also placed in their cluster 11 (our S1), whereas we found this serotype in our S2. The genome they reported (DRR015923) was actually a mixture of *S. boydii* 9 (S1) and *S. boydii* 10 (S2). A recent preprint paper was more comprehensive and used a selection of 485 *Shigella* genomes, selected from over 17,000 publicly available genomes based on ribosomal MLST and ST types. An undescribed phylogeny identified three *Shigella* clusters and seven outliers[15]. The three clusters and five outliers were similar to those from our study. The two remaining, still assigned to *Shigella*, were clusters CSB13 and CSB13-atypical, actually corresponding to *E. albertii*, and attaching and effacing (A/E) *E. coli*, respectively. The status of the provisional serotypes was, however, not addressed adequately, and the continual reporting of different provisional or novel serotypes that are actually identical creates unnecessary confusion for the international surveillance of *Shigella* infections. This issue has now been dealt with by our work (see section "Updating the *Shigella* typing scheme").

**Genomic analysis of the phenotypic markers used in the current *Shigella* typing scheme**

The finding that the O-antigen gene clusters from the *Shigella* serotypes (with the notable exception of *S. sonnei*) were identical to, similar to or adapted from *E. coli* O-antigen gene clusters has been reported before and accounts for the many cross-agglutinations observed with the serotyping scheme[16]. We show here that similarity between *E. coli* and *Shigella rfb* DNA sequences was also identified in *S. dysenteriae* 8 and O38, and in *S. boydii* 22 (formerly *S. boydii* prov. E1621-54) and O7 (Supplementary Fig. 3). For both these *Shigella* serotypes, cross-agglutination with the corresponding *E. coli* serotypes was already reported by Ewing[17]. Finally, the *rfb* from new serotype *S. dysenteriae* 16 (see section "Updating the *Shigella* typing scheme") originated from another species of *Escherichia, E. albertii* (serotype O2).

The molecular basis for some of these metabolic pathways was investigated in previous studies[11,18], and we extend this analysis here to populations not previously considered. An inability to use/ferment mannitol is a key marker of the *S. dysenteriae* serogroup. However, it was also observed in some strains of *S. boydii* 14, *S. flexneri* 4, and *S. flexneri* 6 (ref. [17]). The mannitol (*mtl*) operon contains three genes: *mtlA* encoding the mannitol permease, *mtlD* encoding the mannitol-1-phosphate dehydrogenase, and *mtlR* encoding the mannitol repressor[19,20]. Different mechanisms of mannitol (*mtl*) operon disruption associated with the mannitol-negative trait have occurred independently in *S. dysenteriae*. In the four different subclusters of S1 containing this serotype only remnants of the *mtlA* and *mtlD* genes, flanked by IS*1* and IS*3*, are present in S1a genomes, a remnant of *mtlR* disrupted by one IS*1* is present in S1c genomes, and the entire *mtl* operon is lacking in S1d genomes, represented only by *S. dysenteriae* 7. The analysis of a PacBio sequence of *S. dysenteriae* 7 strain ATCC 9052 (ref. [21]), not included in our study, showed a large IS-driven deletion of the entire operon. The only *S. dysenteriae* in S1b, a historical *S. dysenteriae* 3 strain (Polska 64-3840), was mannitol-positive, with a complete *mtl* operon. We checked the O-antigen agglutination and *in silico* serotyping data for this strain and confirmed its serotype. In the single S2 subcluster containing *S. dysenteriae*, S2d, *mtlR* and, a remnant of *mtlD* were present at a chromosomal location different from that in *E. coli* K-12. The mannitol operon was entirely lacking in the three outlier groups of *S. dysenteriae* (SD1, SD8, and SD10). The precise mechanisms of *mtl* absence remained unclear because the genes flanking *mtl* in *E. coli* K-12 were located >150 kb apart in these three groups.

All strains from the S2 cluster were indole-positive, whereas those from S1 were indole-negative, with one exception, *S. dysenteriae* 7 (indole-positive), assigned to S1d. The loss of indole production has been reported to be due to insertion sequence-mediated insertions and/or deletions damaging the tryptophanase (*tna*) operon in a limited number of strains[18]. This operon, which is 3,144 bp long in *E. coli* K-12, contains three genes: *tnaA* encoding a tryptophanase, *tnaB* encoding a permease, and *tnaL* encoding a 25-residue leader peptide[18]. We confirm here that the loss of indole production is associated with the insertion of an IS*1* at base 55 of *tnaL* in the genomes from subclusters S1a, S1b and S1c. In subcluster S1e, containing rare *S. flexneri* 6 genomes, the *tna* alteration occurred independently, but also involved IS*1* integration, this time in the promoter of *tna*. Additional damage (deletions, other IS insertions) was also observed in individual genomes from these subclusters.

#### Aerogenic strains of *S. boydii* 14 and *S. dysenteriae* 3

By definition, *Shigella* strains do not produce gas from carbohydrate fermentation (except for *S. flexneri* 6), and *S. dysenteriae* strains do not ferment mannitol. The status of aerogenic strains of *S. boydii* 14 long remained a matter of debate. First described in 1943 by Sachs, an aerogenic and mannitol-negative strain referred to as "Enterobacterium A12"[22], was considered, by several authors, to be a variant of *S. boydii* 14 (refs. [23–25]), but was considered by Ewing[26,27] to be an *E. coli* O32. We considered two such aerogenic isolates, including the original Sachs strain A12 (CIP 53.44), in the reference dataset. They were grouped with typical *S. boydii* 14 genomes in the S1b subcluster (including, in particular, the reference strain of this serotype, 2770-51). All five recent *S. boydii* 14 isolates from our routine surveillance dataset were also atypical, as they did not ferment mannitol, and three were aerogenic. Another atypical biotype was described for *S. dysenteriae* 3. Two of these aerogenic and mannitol-positive *S. dysenteriae* 3 strains were described in Poland[28], and the UK[13]. We studied the first of these strains (Polska 64-3840). Unlike aerogenic *S. boydii* 14, strain Polska 64-3840 was not placed by cgMLST in the S1a subcluster with other *S. dysenteriae* 3, but in S2b, with *S. flexneri* 6, which is known to produce gas and to ferment mannitol.

#### Updating the *Shigella* typing scheme

*S. dysenteriae* prov. 93-119 (ref. [29]), prov. SH-103 (ref. [30]), prov. 204/96 (ref. [31]), prov. 96-3162 (ref. [32]), prov. 97-10607 (ref. [33]), prov. SH-105 (referred to as *S. dysenteriae* 16 by Melito and coworkers[30]) belonged to S1a; *S. boydii* prov. 07-6597 (unpublished) belonged to S1c; *S.*

*dysenteriae* prov. 96-265 (ref. [21]), prov. BEDP 02-5104 (ref. [30]), and prov. E670/74 (ref. [34]), belonged to S2d; and *S. boydii* prov. E1621-54 (ref. [35]), belonged to HC1100_11429 within S3.

The representative strains of *S. dysenteriae* prov. 204/96 (# 96-204), prov. 96-3162 (# CDC 96-3162), prov. 97-10607 (# 97-10607), and prov. SH-105 (# CDC97026846) had the same biochemical profile[33], and were all agglutinated with the various antisera raised against strains 96-204, 97-10607, SH-105, and KIVI 162. This last strain was not included in this study, but was reported as a new *S. dysenteriae* serotype in Bangladesh[36]. The representative strains of *S. dysenteriae* serotypes prov. 204/96, prov. 96-3162, prov. 97-10607, and prov. SH-105 belonged to HC100_44952 and had identical complete *rfb* clusters of 17·3 kb in size and displaying 100% identity (17,306/17,306), with no gaps relative to the *rfb* cluster of *E. albertii* O2 strain SP140152 (GenBank accession code KY574563)[37]. A partial match (91% identity, 7,041/7,744 with three gaps) was also obtained with the *rfb* of *S. dysenteriae* 4 (CP026840). This 17·3 kb *rfb* cluster was not the normal *rfb* cluster (i.e., that located next to the colanic acid biosynthesis gene cluster). Instead, it consisted only of a remnant of the *S. dysenteriae* 3 *rfb* cluster, damaged by IS*5* (Supplementary Fig. 4). The 17·3 kb *rfb* cluster was located on a ~63 kb genomic island integrated close to the *yqjH* gene (Supplementary Fig. 5). Based on *rfb* sequences, 123 of the 133 HC100_44952 genomes in EnteroBase (6 May 2021) belong to this provisional *S. dysenteriae* serotype, and seven are genuine *S. dysenteriae* 3. The new serotype of *S. dysenteriae* described in the UK as E112707/96 was not included in our study[38]. However, an analysis of its representative genome deposited in EnteroBase (SRR4195641) revealed that it had the same 17·3 kb *rfb* cluster as strain 96-204 and also belonged to HC100_44952. Thus, this novel serotype, described by different groups across the world under different names, is actually identical for all strains considered. It has also become relatively frequent. It ranked first and accounted for 30·2% (16/53) of the *S. dysenteriae* isolates in our French routine surveillance dataset (2017-2020). Between 2004 and 2017, 19% (150/754) of the *S. dysenteriae* isolates received by Public Health England also belonged to E112707/96 (ref. [38]). We therefore propose its addition to the *Shigella* typing scheme under the name *S. dysenteriae* 16. Strain 96-204 has been deposited in the *Collection de l'Institut Pasteur* (CIP) as the reference strain for this serotype, under CIP 111935.

The representative strains of *S. dysenteriae* prov. 93-119 (# 93-119) and prov. SH-103 (# CDC95011241) had identical biochemical profiles (Supplementary Table 7) and both were agglutinated with an antiserum raised against strain 93-119. They clustered into HC100_35368 and had identical 16·3 kb *rfb* clusters (Supplementary Fig. 6). This *rfb* cluster displayed no similarity to those of *Shigella* genomes, but was similar (identity:16,307/16,323; gap, 1/16,323) to that of *E. coli* strain YSP-8 (GenBank accession no. CP037910), a non-serotyped *E. coli* collected from pig faeces in China. On the basis of their *rfb* sequences, 17 of the 20 HC100_35368 isolates present in EnteroBase in May 2021 belonged to this new serotype and three were genuine *S. dysenteriae* 6. We propose to add this new serotype to the *Shigella* serotyping scheme under the name *S. dysenteriae* 17. Strain 93-119 has been deposited in the CIP as the reference strain for this serotype, under CIP 111948.

The three strains from *S. boydii* prov. 07-6597 identified here, associated with travel to Morocco, Chad and Uzbekistan, clustered within S1c, close to the genomes of *S. boydii* 1 and 20 (Fig. 2). The three *S. boydii* prov. 07-6597 strains were agglutinated with a serum against strain 07-6597 developed in-house. Their biochemical profile is shown in Supplementary Table 7. The three strains belonged to HC50_45442 and had identical *rfb* clusters of 17·2 kb in size (Supplementary Fig. 6) that were similar (identities, 17,135/17,260, 99%; gap, 1/17,620) to the *rfb* cluster of *E. coli* O180 strain 86-381 (GenBank accession code AB812077) (Supplementary

Fig. 3). Interestingly, the first 4 kb of the *S. boydii* prov. 07-6597 *rfb* partially matched that of *S. boydii* 1 (identities 4,230/4,644, 91%; gaps 3/4,644). A search of EnteroBase (6 May 2021) found 16 additional *S. boydii* prov. 07-6597 genomes, all in HC50_45442 (two *S. boydii* 1 genomes were also found in HC50_45442). We therefore propose to add this new serotype to the *Shigella* serotyping scheme under the name *S. boydii* 21. Strain 07-6597 has been deposited in the CIP as the reference strain for this serotype, under CIP 111949.

A non-serotypable *S. boydii* strain (# 07-7164) clustered within S1c, close to genomes of *S. boydii* 1 and 20 (Fig. 2). The *rfb* cluster of 07-7164 was similar to that of *S. boydii* 20 (itself derived from that of *S. boydii* 1 via one or two IS*1* insertions, Supplementary Fig. 6), except that 07-7164 carried an additional IS*1*, inserted within the *wzy* gene and associated with a deletion encompassing the *wbuU* and *wbuW* genes.

*S. dysenteriae* prov. BEDP 02-5104 (also known as prov. SH-111)[30], and prov. 96-265 clustered together in S2d. Both had two *rfb* clusters: one chromosomal (at the normal site close to the colanic gene cluster) and 10·8 kb in size, and the other plasmid-borne and 11·9 kb in size. The chromosomal cluster was identical to that of *S. dysenteriae* 2, whereas the plasmid-borne cluster was similar (99-100% identities with 11,981/11,981 or 11,978/11,981; no gaps) to the plasmid-borne *rfb* clusters of *E. coli* and *Citrobacter* (GenBank accessions nos. CP048012 and AP022515). These isolates were agglutinated with a serum raised against BEDP 02-5104 but not by the anti-*S. dysenteriae* 2 typing serum, suggesting that expression of the plasmid-borne *rfb* genes has superseded the expression of the chromosomal genes. The plasmid also carried a raffinose operon, accounting for the use of this trisaccharide by *S. dysenteriae* prov. BEDP 02-5104, a very unusual trait in *S. dysenteriae*[30]. In EnteroBase, 45 of the 46 genomes with both *rfb* clusters belonged to HC100_11651, which consisted exclusively of these *S. dysenteriae* prov. BEDP 02-5104 genomes (Supplementary Fig. 7). The remaining genome was located in another HC100 cluster, among *S. dysenteriae* 2 genomes. This analysis suggests that this provisional serotype is actually a *S. dysenteriae* 2 that has acquired an O-antigen-modifying plasmid. In the absence of knowledge about the stability of this plasmid and the possibility of its transfer to *S. dysenteriae* 2 from other HC100 clusters, it seems wise not to consider *S. dysenteriae* prov. BEDP 02-5104 to be a new serotype for the time being.

The representative strain of *S. dysenteriae* prov. E670/74 (# E670/74) was also found in S2d, along with *S. dysenteriae* 2. Its *rfb* cluster was 9·7 kb in size and was similar (99% identities, 9,646/9,709; gaps, 0/9,709) to that of *E. coli* O170 (GenBank accession no. AB812070.1)[39] (Supplementary Figs. 3 and 6). This provisional serotype was described in 1989 and has not since been reported[34]. We did not identify it in our routine surveillance dataset either. Only four of the 222 S2d genomes in EnteroBase in May 2021 had this 9·7 kb *rfb* cluster, including at least three independent cultures of the same strain, E670/74. Should this serotype be isolated sporadically, it would be considered non-typable in the absence of a dedicated typing serum and genomic sequencing, and would therefore remain undetected. We, therefore, suggest its addition to the *Shigella* serotyping scheme under the name *S. dysenteriae* 18. Strain E670/74 was deposited in the National Collection of Type Cultures (NCTC) as the reference strain for this serotype, under NCTC 11311.

The three *S. boydii* prov. E1621-54 strains studied[13,35], including # E1621-54, belonged to a particular cluster (HC1100_11429) of S3. All the other S3 isolates were *S. flexneri* (serotypes X, Y, and 1-5). These *S. boydii* prov. E1621-54 genomes had an identical 16·9 kb *rfb* cluster, similar (99% identities, 16,879/16,919; gaps, 2/16,919) to that of *E. coli* O7:K1 (GenBank accession no. CP003034) (Supplementary Figs. 3 and 6). This *rfb* was more distantly related to that of *S. boydii* 12 (Supplementary Fig. 6). The strains of this provisional serotype – originally described in humans and monkeys from Indonesia – produced indole[13,35]. However, most of our *S. boydii* prov. E1621-54 isolates were indole-negative. This loss of function was associated with an IS*1* inserted into the promoter region of the *tna* operon (different from the insertion in S1e). One additional *S. boydii* prov. E1621-54 isolate was identified in our routine surveillance dataset. In EnteroBase, approximately 60 non-redundant genomes belonged to HC1100_11429, and all had the 16·9 kb *rfb* cluster of *S. boydii* prov. E1621-54. We therefore propose the addition of this serotype to the *Shigella* typing scheme under the name *S. boydii* 22. Strain E1621-54 has been deposited in the CIP as the reference strain for this serotype, under CIP 111950.

**Supplementary Table 1.** cgMLST allelic differences between the PacBio and Illumina genomes obtained for identical strains.

| PacBio genome | Illumina genome | Serotype[1] | cgMLST allelic distances |
|---|---|---|---|
| **54/1621** | E1621-54 | SB 22 | 288 |
| **ATCC 49812** | CIP 57.47 | SB 9 | 67 |
| **ATCC 13313** | CIP 57.28 | SD 1 | 29 |
| **ATCC 49346** | E22383 | SD 14 | 163 |
| **ATCC 49347** | E23507 | SD 15 | 48 |
| **96-3162** | 96-3162 | SD 16 | 365 |
| **204/96** | 96-204 | SD 16 | 9 |
| **E670/74** | E670/74 | SD 18 | 24 |
| **ATCC 9754** | CIP 52.32 | SD 6 | 584 |
| **ATCC 12037** | CIP 58.26 | SD 9 | 12 |
| **74-1170** | NCDC1170-74 | SF 5a | 16 |

[1]SB, *S. boydii*; SD, *S. dysenteriae*; SF, *S. flexneri*

**Supplementary Table 2.** GenBank accession numbers of the *Shigella* O-antigen gene clusters.

| O-antigen gene cluster from: | Accession number |
|---|---|
| *S. dysenteriae* 1 | MZ286368 |
| *S. dysenteriae* 2 | MZ286369 |
| *S. dysenteriae* 3 | MZ286370 |
| *S. dysenteriae* 4 | MZ286371 |
| *S. dysenteriae* 5 | MZ286372 |
| *S. dysenteriae* 6 | MZ286373 |
| *S. dysenteriae* 7 | MZ286374 |
| *S. dysenteriae* 8 | MZ286375 |
| *S. dysenteriae* 9 | MZ286376 |
| *S. dysenteriae* 10 | MZ286364 |
| *S. dysenteriae* 11 | MZ286365 |
| *S. dysenteriae* 12 | MZ286366 |
| *S. dysenteriae* 13 | MZ286367 |
| *S. dysenteriae* 14 | MF322749 |
| *S. dysenteriae* 15 | MF322748 |
| *S. dysenteriae* 16 (prov. 204/96) | MF322751 |
| *S. dysenteriae* 17 (prov. 93-119) | MF322752 |
| *S. dysenteriae* 18 (prov. E670/74) | MF322750 |
| *S. dysenteriae* prov. BEDP 02-5104 | MZ303046 |
| *S. boydii* 1 | MZ286385 |
| *S. boydii* 2 | MZ286387 |
| *S. boydii* 3 | MZ286388 |
| *S. boydii* 4 | MZ286389 |
| *S. boydii* 5 | MZ286390 |
| *S. boydii* 6 | AF402314 |
| *S. boydii* 7 | MZ286391 |
| *S. boydii* 8 | MZ286392 |
| *S. boydii* 9 | MZ286393 |
| *S. boydii* 10 | MZ286378 |
| *S. boydii* 11 | MZ286379 |
| *S. boydii* 12 | EU296406 |
| *S. boydii* 14 | MZ286380 |
| *S. boydii* 15 | MZ286381 |
| *S. boydii* 16 | MZ286382 |
| *S. boydii* 17 | MZ286383 |
| *S. boydii* 18 | MZ286384 |
| *S. boydii* 19 | MF322754 |
| *S. boydii* 20 | MZ286386 |
| *S. boydii* 21 (prov. 07-6597) | MF322754 |
| *S. boydii* 22 (prov. E1621-54) | MF322747 |
| *S. flexneri* 1-5, X, Y | MZ286377 |
| *S. flexneri* 6 | MZ286394 |
| *S. sonnei* (plasmid) | AF285971 |
| *S. sonnei* (chromosome) | AF031957 |

**Supplementary Table 3.** GenBank accession numbers and coordinates of the gene and genetic structures studied.

| Target | Strain | Accession no. | Coordinates |
|---|---|---|---|
| ***mtlA*** | *Escherichia coli* K-12 substr. MG1655 | NC_000913.3 | 3772281-3774194 |
| ***mtlD*** | *Escherichia coli* K-12 substr. MG1655 | NC_000913.3 | 3774424-3775572 |
| ***mtlR*** | *Escherichia coli* K-12 substr. MG1655 | NC_000913.3 | 3775572-3776159 |
| ***tnaC*** | *Escherichia coli* K-12 substr. MG1655 | NC_000913.3 | 3888435-3888509 |
| ***tnaA*** | *Escherichia coli* K-12 substr. MG1655 | NC_000913.3 | 3888730-3890145 |
| ***tnaB*** | *Escherichia coli* K-12 substr. MG1655 | NC_000913.3 | 3890236-3891483 |
| ***ipaH*** | *Shigella boydii* CDC 3083-94 | CP001063.1 | 1918645-1920360 |
| ***rafABD*** | *Escherichia coli* K-12 | M27273.1 | 1-5284 |
| ***rafY*** | *Escherichia coli* | U82290.1 | 1-1866 |

**Supplementary Table 4.** Number of *Shigella* strains and isolates studied per dataset.

| Serotype | Reference | Reference+ | Routine |
|---|---|---|---|
| ***S. boydii*** | **70** | **10** | **101** |
| 1 | 3 | 0 | 5 |
| 2 | 3 | 0 | 40 |
| 3 | 3 | 0 | 0 |
| 4 | 5 | 1 | 10 |
| 5 | 4 | 2 | 2 |
| 6 | 1 | 0 | 0 |
| 7 | 3 | 0 | 0 |
| 8 | 3 | 0 | 2 |
| 9 | 4 | 3 | 1 |
| 10 | 4 | 0 | 7 |
| 11 | 5 | 0 | 14 |
| 12 | 2 | 2 | 0 |
| 14 | 6 | 0 | 5 |
| 15 | 2 | 0 | 0 |
| 16 | 2 | 0 | 0 |
| 17 | 1 | 1 | 0 |
| 18 | 3 | 1 | 4 |
| 19 | 3 | 0 | 5 |
| 20 | 6 | 0 | 5 |
| 21 | 3 | 0 | 0 |
| 22 | 3 | 0 | 1 |
| Rough | 0 | 0 | 0 |
| NST | 1 | 0 | 0 |
| ***S. dysenteriae*** | **81** | **2** | **53** |
| 1 | 16 | 0 | 0 |
| 2 | 7 | 0 | 14 |
| 3 | 3 | 0 | 7 |
| 4 | 4 | 0 | 1 |
| 5 | 3 | 0 | 0 |
| 6 | 2 | 0 | 0 |
| 7 | 3 | 0 | 0 |
| 8 | 3 | 2 | 0 |
| 9 | 3 | 0 | 1 |
| 10 | 2 | 0 | 0 |
| 11 | 2 | 0 | 0 |
| 12 | 4 | 0 | 7 |
| 13 | 2 | 0 | 0 |
| 14 | 1 | 0 | 0 |
| 15 | 1 | 0 | 0 |
| 16 | 6 | 0 | 16 |
| 17 | 6 | 0 | 1 |
| 18 | 1 | 0 | 0 |
| prov. BEDP 02-5104 | 12 | 0 | 2 |
| Rough | 0 | 0 | 3 |
| NST | 0 | 0 | 1 |
| ***S. sonnei*** | **44** | **0** | **2161** |
| Lineage I | 11 | 0 | 14 |
| Lineage II | 15 | 0 | 108 |
| Lineage III | 17 | 0 | 2039 |
| Lineage IV | 1 | 0 | 0 |
| ***S. flexneri* 1-5** | **103** | **65** | **1423** |
| **Lineage I** | **32** | **12** | **460** |
| 1a | 6 | 1 | 9 |
| 1b | 9 | 0 | 338 |
| 1c/7a | 2 | 3 | 74 |
| 7b | 3 | 0 | 11 |
| 2a | 0 | 0 | 1 |

| | | | |
|---|---|---|---|
| 2b | 2 | 2 | 1 |
| 3b | 3 | 0 | 8 |
| 4a | 0 | 2 | 1 |
| 4av | 2 | 3 | 12 |
| 4b | 3 | 1 | 0 |
| 4bv | 0 | 0 | 0 |
| X | 1 | 0 | 1 |
| Y | 1 | 0 | 1 |
| Yv | 0 | 0 | 3 |
| **Lineage II** | **25** | **2** | **218** |
| 3a | 18 | 2 | 200 |
| 3b | 7 | 0 | 16 |
| Unknown | 0 | 0 | 1 |
| X | 0 | 0 | 0 |
| Y | 0 | 0 | 1 |
| **Lineage III** | **31** | **10** | **680** |
| 1a | 1 | 1 | 1 |
| 2 | 0 | 0 | 2 |
| 2a | 19 | 0 | 599 |
| 2b | 1 | 5 | 40 |
| 4 | 0 | 0 | 0 |
| 5a | 2 | 0 | 0 |
| X | 1 | 0 | 2 |
| Xv | 4 | 0 | 20 |
| Y | 1 | 4 | 14 |
| Yv | 2 | 0 | 1 |
| Rough | 0 | 0 | 1 |
| **Lineage IV** | **9** | **9** | **11** |
| 3a | 5 | 1 | 11 |
| 3b | 1 | 2 | 0 |
| 4bv | 1 | 1 | 0 |
| X | 2 | 1 | 0 |
| NST | 0 | 4 | 0 |
| **Lineage V** | **2** | **8** | **0** |
| 5a | 2 | 4 | 0 |
| 5b | 0 | 4 | 0 |
| **Lineage VI** | **2** | **14** | **3** |
| Y | 2 | 5 | 0 |
| Yv | 0 | 7 | 3 |
| Xv | 0 | 2 | 0 |
| **Lineage VII** | **2** | **10** | **51** |
| 4 | 0 | 0 | 1 |
| 4a | 0 | 1 | 12 |
| 4av | 1 | 3 | 31 |
| Y | 0 | 5 | 0 |
| Yv | 1 | 1 | 7 |
| ***S. flexneri* 6** | **19** | **4** | **132** |
| Boyd 88 | 12 | 0 | 120 |
| Hertfordshire | 4 | 0 | 9 |
| Manchester | 2 | 0 | 3 |
| Newcastle | 1 | 0 | 0 |
| Unknown | 0 | 4 | 0 |
| **TOTAL** | **317** | **81** | **3870** |

NST, non-serotypable

**Supplementary Table 5.** Distribution of the different *Shigella* serotypes in clusters S1 and S2 of 398 *Shigella* reference and reference+ genomes according to their HC2000, HC1100, and HC400 data.

| **Cluster** | **HC2000** | **Subcluster** | **HC1100** | **HC400** | **Serotype[1]** |
|---|---|---|---|---|---|
| **S1** | 1465 | S1a | 4194 | 4194 | SD3, SD13, SD15 |
| | | | | 14114 | SD11, SD12 |
| | | | | 17375 | SD4 |
| | | | | 35330 | SD3, SD12, SD14, SD16 |
| | | | | 35368 | SD6, SD17 |
| | | | | 44956 | SD9 |
| | | | | 45269 | SD9 |
| | | | | 45271 | SD11 |
| | | S1b | 1465 | 1465 | SB2, SB4, SB11 |
| | | | | 11126 | SF6 |
| | | | | 11341 | SF6 |
| | | | | 13048 | SF6 |
| | | | | 17342 | SF6 |
| | | | | 22378 | SD3, SF6 |
| | | | | 41808 | SB14 |
| | | | | 45451 | SF6 |
| | | S1c | 1466 | 1466 | SB1, SB8, SB18, SB19, SB20, SB21 |
| | | | | 45284 | SD5 |
| | | | | 45300 | SB6, SB10 |
| | | | | 45420 | SB3 |
| | | S1d | 36524 | 36524 | SD7 |
| | | S1e | 45518 | 45518 | SF6 |
| **S2** | 4118 | S2a | 11452 | 11452 | SB17 |
| | | | | 11601 | SB5 |
| | | S2b | 4118 | 4118 | SB5 |
| | | | | 11449 | SB16 |
| | | S2c | 11421 | 11421 | SB11 |
| | | S2d | 4191 | 4191 | SD2, SD18 |
| | | | | 11444 | SB15 |
| | | | | 11651 | SD2, SD prov. BEDP 02-5104 |
| | | | | 44479 | SD2 |
| | | S2e | 7057 | 11413 | SB9 |
| | | | | 11414 | SB9 |
| | | | | 30095 | SB9 |
| | | | | 61169 | SB9 |
| | | S2f | 11401 | 11401 | SB7 |

[1]SB, *S. boydii*; SD, *S. dysenteriae*; SF, *S. flexneri*

**Supplementary Table 6.** *In silico* serotype prediction for 316 *Shigella* reference strains with serotype designation, obtained with various tools.

| | | % assignment: | | | | | | | | | | | | | | | |
|---|---|---|---|---|---|---|---|---|---|---|---|---|---|---|---|---|---|
| | | **EnteroBase SeroPred** | | | | **ShigaTyper** | | | | **ShigEiFinder (Fasta)** | | | | **ShigEiFinder (Reads)** | | | |
| **Serotype** | *n* | C | U | I | N | C | U | I | N[#] | C | U | I | N | C | U | I | N |
| ***S. boydii*** | | | | | | | | | | | | | | | | | |
| **1** | 3 | 0 | 100 (B1/O149) | | | 100 | | | | 100 | | | | 100 | | | |
| **2** | 3 | 100 | | | | 67 | | | 33 | 100 | | | | 100 | | | |
| **3** | 3 | 100 | | | | 100 | | | | 100 | | | | 100 | | | |
| **4** | 5 | 20 | 80 (B4/O53) | | | 80 | | | 20 | 100 | | | | 100 | | | |
| **5** | 4 | 100 | | | | 50 | | | 50 | 100 | | | | 100 | | | |
| **6** | 1 | 100 | | | | | | | 100 | 100 | | | | 100 | | | |
| **7** | 3 | 100 | | | | 100 | | | | 100 | | | | 100 | | | |
| **8** | 3 | 100 | | | | 100 | | | | 100 | | | | 100 | | | |
| **9** | 4 | 100 | | | | 100 | | | | 100 | | | | 100 | | | |
| **10** | 4 | | | 100 (B6) | | | | | 100 | | | 100 (B6) | | | | 100 (B6) | |
| **11** | 5 | 60 | | 20 (O105) | 20 | 60 | | 20 (Not) | 20 | 80 | 20 (Cs1) | | | 60 | 20 (Cs1) | 20[!] | |
| **12** | 2 | 100 | | | | 100 | | | | 100 | | | | 100 | | | |
| **14** | 6 | 100 | | | | 100 | | | | 83 | 17 (Cs1) | | | 100 | | | |
| **15** | 2 | | 100 (B15/O112) | | | 100 | | | | 100 | | | | 100 | | | |
| **16** | 2 | 100 | | | | 100 | | | | 100 | | | | 100 | | | |
| **17** | 1 | 100 | | | | 100 | | | | 100 | | | | 100 | | | |
| **18** | 3 | 100 | | | | 100 | | | | 100 | | | | 100 | | | |
| **19** | 3 | | | | 100 | 67 | | | 33 | 100 | | | | 100 | | | |
| **20** | 6 | | | 100 (B1/O149) | | 67 | | | 33 | 100 | | | | 100 | | | |
| **21*** | 3 | | | 100 (O180) | | | | 67 (B18) | 33 | | 100 (Cs1/O180) | | | | 100 (Cs1/O180) | | |
| **22*** | 3 | | | 100 (O7) | | 100 | | | | 100 | | | | 100 | | | |
| ***S. dysenteriae*** | | | | | | | | | | | | | | | | | |
| **1** | 16 | 100 | | | | 100 | | | | 100 | | | | 100 | | | |
| **2** | 7 | 86 | | | 14 | 100 | | | | 86 | 14 (Cs2) | | | 86 | 14 (Cs2) | | |
| **3** | 3 | | | 100 (O124, O164) | | 100 | | | | 100 | | | | 100 | | | |
| **4** | 4 | | | 75 (O168/OX6) | 25 | 100 | | | | 100 | | | | 75 | 25 (Cs1) | | |
| **5** | 3 | | | 100 (O58) | | 100 | | | | 100 | | | | 100 | | | |
| **6** | 2 | | | 100 (O130) | | 100 | | | | 100 | | | | 100 | | | |

| | | | | | | | | | | | | | | | | |
|---|---|---|---|---|---|---|---|---|---|---|---|---|---|---|---|---|
| **7** | 3 | | | 100 (O121) | | 100 | | | | 100 | | | | 100 | | |
| **8** | 3 | | | 100 (O38) | | 100 | | | | 100 | | | | 100 | | |
| **9** | 3 | | | 100 (O40) | | 100 | | | | 100 | | | | 100 | | |
| **10** | 2 | 50 | | | 50 | 100 | | | | 100 | | | | 100 | | |
| **11** | 2 | | 50 (D11/O29) | 50 (O29) | | 100 | | | | 100 | | | | 100 | | |
| **12** | 4 | | | 100 (O152) | | 100 | | | | 100 | | | | 100 | | |
| **13** | 2 | | | 100 (O150) | | 100 | | | | 100 | | | | 100 | | |
| **14** | 1 | | | | 100 | 100 | | | | 100 | | | | 100 | | |
| **15** | 1 | | | | 100 | 100 | | | | 100 | | | | 100 | | |
| **16*** | 6 | | | | 100 | | | 67 (p. 96-265) | 33 | | 17 (Cs1) | 83 (p. 96-265) | | | 17 (Cs1) | 83 (p. 96-265) |
| **17*** | 6 | | | | 100 | | | | 100 | | 100 (Cs1) | | | | 100 (Cs1) | |
| **18*** | 1 | | | 100 (O170) | | 100 | | | | 100 | | | | 100 | | |
| **p. 02-5104** | 12 | | | 100 (D2) | | | | 100 (D2) | | | | 100 (D2) | | | | 100 (D2) |
| ***S. flexneri*** | | | | | | | | | | | | | | | | |
| **1-5, X, Y** | 10 3 | 58 | 36 (F1-5/O13) | 1 (O129/O13) | 5 | 100 | | | | 100 | | | | 100 | | |
| **1a** | 7 | | | | 100 | 100 | | | | 100 | | | | 100 | | |
| **1b** | 9 | | | | 100 | 100 | | | | 100 | | | | 100 | | |
| **1c** | 2 | | | | 100 | 100 | | | | | 100 (1c/7b) | | | | 100 (1c/7b) | |
| **2a** | 19 | | | | 100 | 100 | | | | 100 | | | | 100 | | |
| **2b** | 3 | | | | 100 | 100 | | | | 100 | | | | 100 | | |
| **3a** | 23 | | | | 100 | 100 | | | | 100 | | | | 100 | | |
| **3b** | 11 | | | | 100 | 73 | 18 (new) | 9 (1b) | | 73 | 18[¥] | 9 (1b) | | 73 | 18[¥] | 9 (1b) |
| **4av** | 3 | | | | 100 | 67 | | 33 (4bv) | | 67 | 33 (4av/4b) | | | 67 | 33 (4av/4b) | |
| **4b** | 3 | | | | 100 | 100 | | | | 100 | | | | 100 | | |
| **4bv** | 1 | | | | 100 | 100 | | | | | 100 (4av/4b) | | | | 100 (4av/4b) | |
| **5a** | 4 | | | | 100 | 100 | | | | 100 | | | | 100 | | |
| **7b** | 3 | | | | 100 | 100 | | | | | 100[¥] | | | | 100[¥] | |
| **X** | 4 | | | | 100 | 50 | | 50 (3a, Xv) | | 75 | | 25 (3a) | | 75 | | 25 (3a) |
| **Xv** | 4 | | | | 100 | 100 | | | | 100 | | | | 100 | | |
| **Y** | 4 | | | | 100 | 50 | | 50 (X) | | 50 | | 50 (X) | | 50 | | 50 (X) |
| **Yv** | 3 | | | | 100 | 67 | | 33 (Xv) | | 100 | | | | 100 | | |
| **6** | 19 | 79 | | 21 (O147) | | 95 | | | 5 | 95 | | 5[!] | | 100 | | |
| ***S. sonnei*** | 44 | | | | 100 | 75 | | | 25 | 95 | | 5[!] | | 97.7 | | 2.3[!] |

C, correct; U, uncertain; I, incorrect; N, none; *, novel *Shigella* serotype described in our study; #, no prediction for ShigaTyper (no *wzx*, multiple *wzx* or error); new, novel serotype; Not, not *Shigella* or EIEC; [!], *Shigella* or EIEC unclustered; B, *S. boydii*; D, *S. dysenteriae*; F, *S. flexneri*; p. 96-265, *S. dysenteriae* prov. 96-265; Cs1, Cluster 1 (*Shigella*) from ref. [15] ; Cs2, Cluster 2 (*Shigella*) from ref. [15] ; [¥], Cluster 3 (*S. flexneri*) from ref. [15]

**Supplementary Table 7.** Biochemical characteristics of new serotypes *S. dysenteriae* 17 and *S. boydi* 21.

| ***Test*** | *S. dysenteriae 17* | *S. boydii 21* |
|---|---|---|
| Motility | - | - |
| Oxidase | - | - |
| β-galactosidase (ONPG) | - | - |
| Lysine decarboxylase | - | - |
| Ornithine decarboxylase | - | - |
| Arginine dihydrolase | - | - |
| Tryptophan deaminase | - | - |
| Indole production | - | - |
| Voges-Proskauer (37°C) | - | - |
| Urea hydrolysis | - | - |
| $H_2S$ production | - | - |
| Gelatinase | - | - |
| $NO_2$ from $NO_3$ | + | + |
| Citrate utilization | - | - |
| Glucose (gas) | - | - |
| Acid from: | | |
| D-adonitol | - | - |
| Amidon | - | - |
| L-arabinose | -[a] | + |
| D-cellobiose | - | - |
| Dulcitol | - | - |
| D-fructose | + | + |
| L-fucose | - | - |
| D-galactose | + | + |
| D-glucose | + | + |
| Glycerol | (+) | (+) |
| Inositol | - | - |
| D-lactose | - | - |
| D-maltose | - | - |
| D-mannitol | - | + |
| D-mannose | + | + |
| N-acetyl glucosamine | + | + |
| Potassium 2-keto-gluconate | - | - |
| Potassium 5-keto-gluconate | - | - |
| Potassium gluconate | + | + |
| D-raffinose | - | - |
| L-rhamnose | - | - |
| L-ribose | + | + |
| Salicin | - | - |
| D-sorbitol | -[b] | -[c] |
| L-sorbose | (+) | - |
| D-sucrose | - | - |
| D-trehalose | + | + |
| D-xylose | - | - |
| L-xylose | - | - |

+, all strains positive (one day); -, all strains negative; (+), all strains positive (two days); [a], positive in one day with the API 50 CH strip (negative with the API 20 E strip); [b], negative or positive in two days with the API 50 CH strip (negative with the API 20 E strip); [c], positive in two days with the API 50 CH strip (negative with the API 20 E strip).

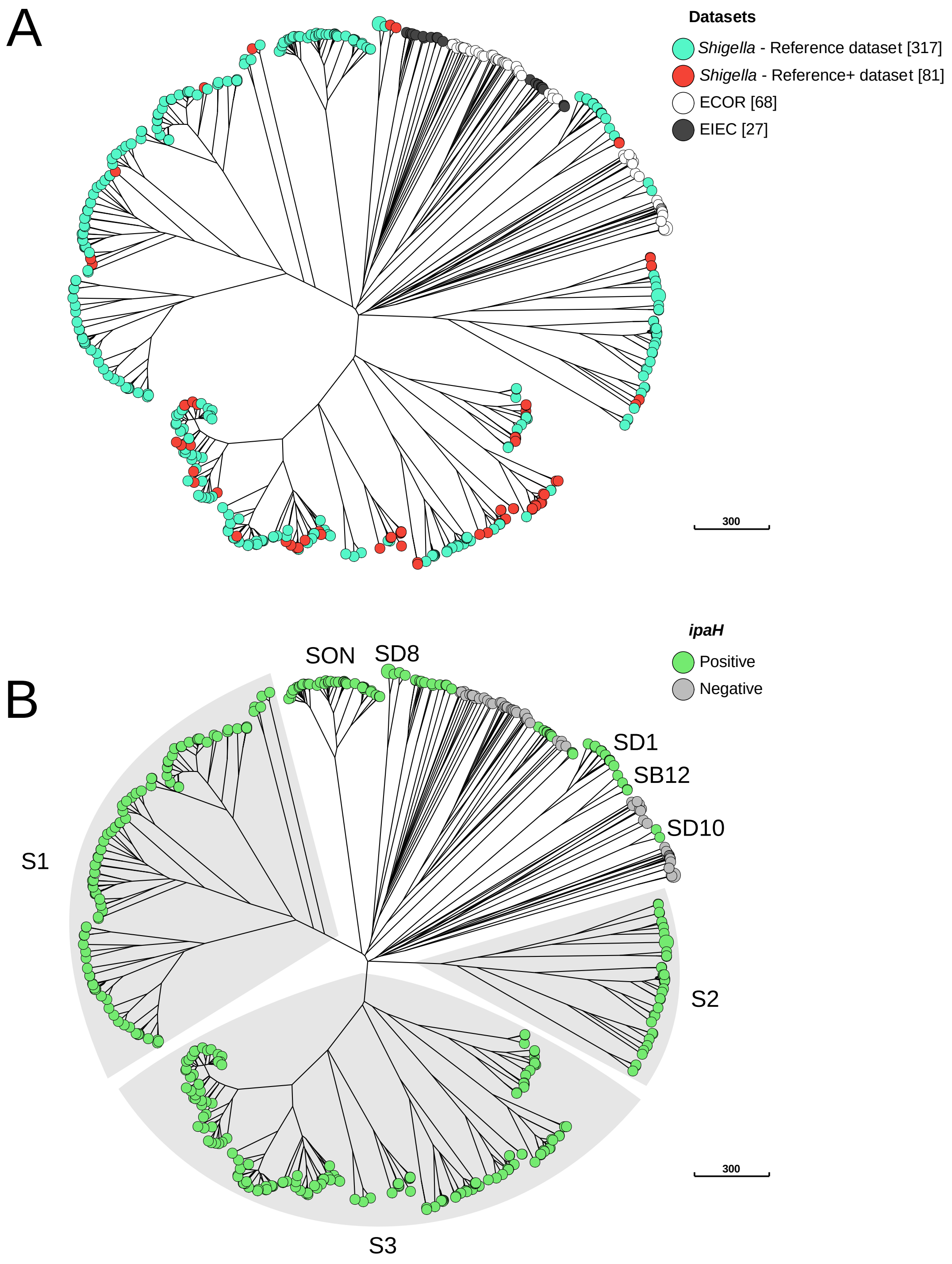

**Supplementary Figure 1**. A NINJA neighbour-joining GrapeTree showing the population structure of *Shigella* spp. based on the cgMLST allelic differences between 493 *Shigella* and *E. coli* reference genomes. This tree is the same as that shown in Figure 1. (A) The tree nodes are colour-coded by type of dataset, to illustrate the differential contribution to *Shigella* population diversity of the reference and reference+ datasets. (B) The tree nodes are colour-coded according to the presence or absence of the *ipaH* gene. The numbers of isolates in each dataset are indicated in brackets.

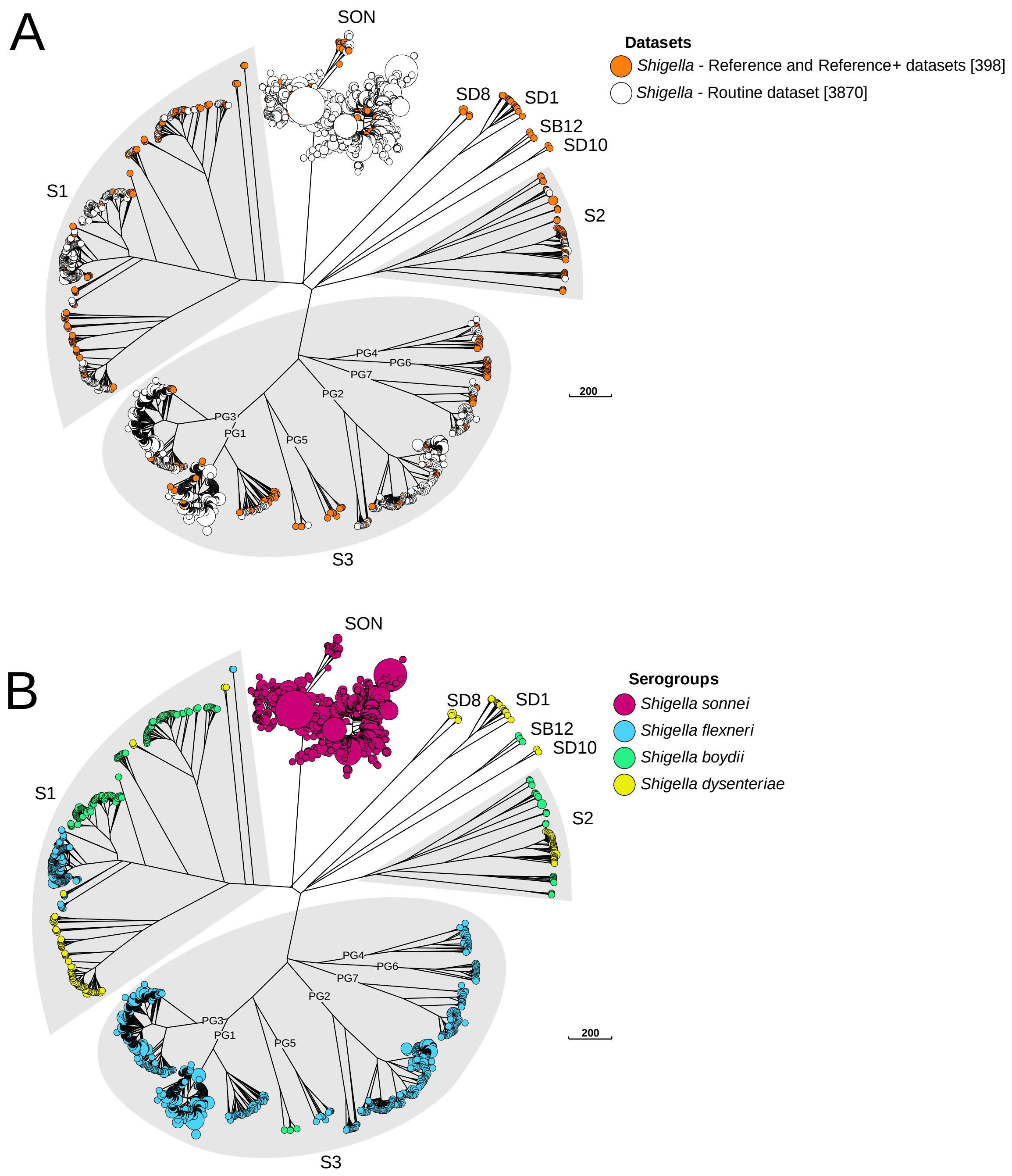

**Supplementary Figure 2.** A NINJA neighbour-joining GrapeTree showing the genomic diversity of 4,268 *Shigella* isolates, based on the reference, reference+ and routine surveillance datasets. The tree nodes are colour-coded by type of dataset (A) or by *Shigella* serogroup (B). The global dataset consists of the reference and reference+ datasets. The numbers of isolates in each dataset are indicated in brackets.

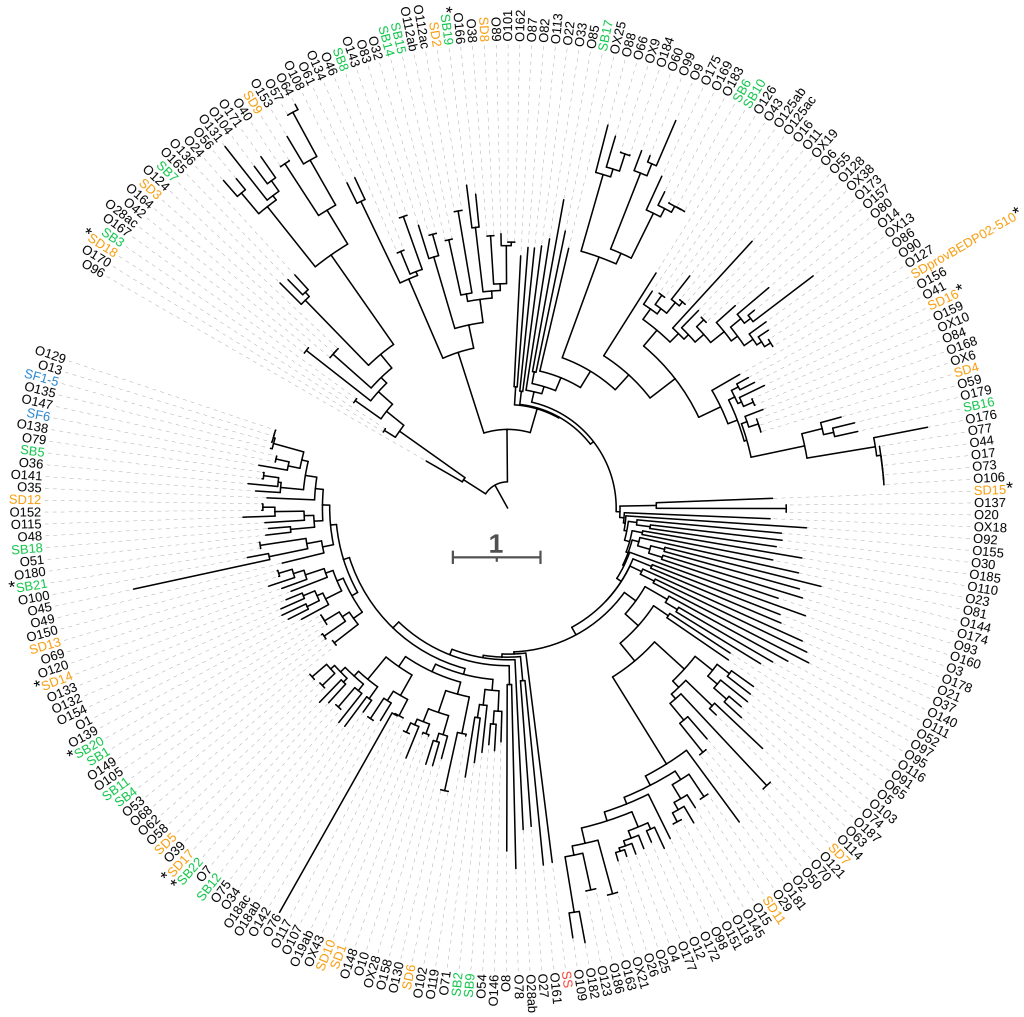

**Supplementary Figure 3.** Phylogenetic tree for all *Shigella* and *E. coli* O-antigen gene clusters. The *Shigella* O-antigen gene clusters (*rfb*) characterised in this study are indicated by an asterisk. Nodes supported by bootstrap values ≥95% are indicated by red dots. *S. boydii rfb* genes are shown in green, S. *dysenteriae rfb* genes are shown in orange, *S. flexneri rfb* genes are shown in blue, and *S. sonnei rfb* genes are shown in red.

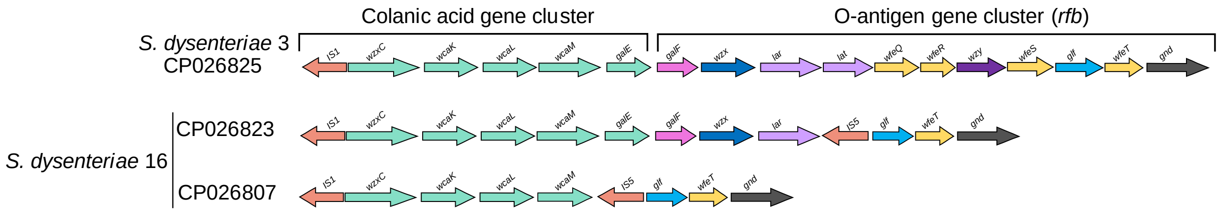

**Supplementary Figure 4:** Representation of the normal O-antigen gene cluster locus in *S. dysenteriae* 3 and *S. dysenteriae* 16 isolates.

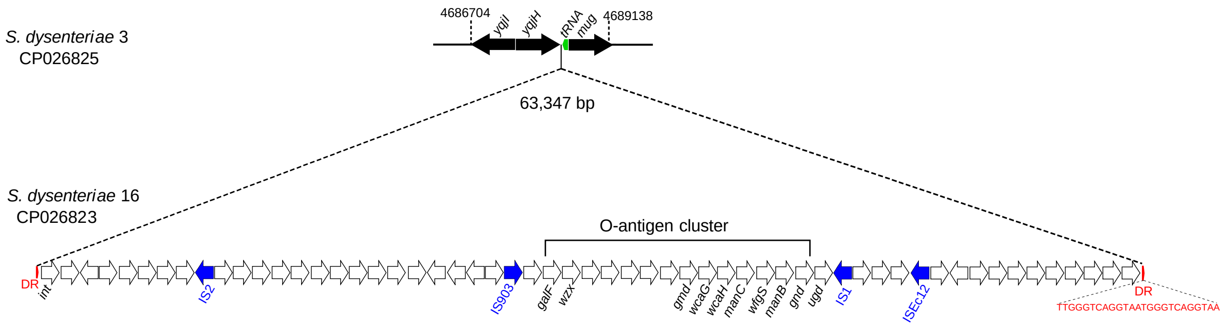

**Supplementary Figure 5:** Representation of the genomic island containing the complete O-antigen cluster in *S. dysenteriae* 16.

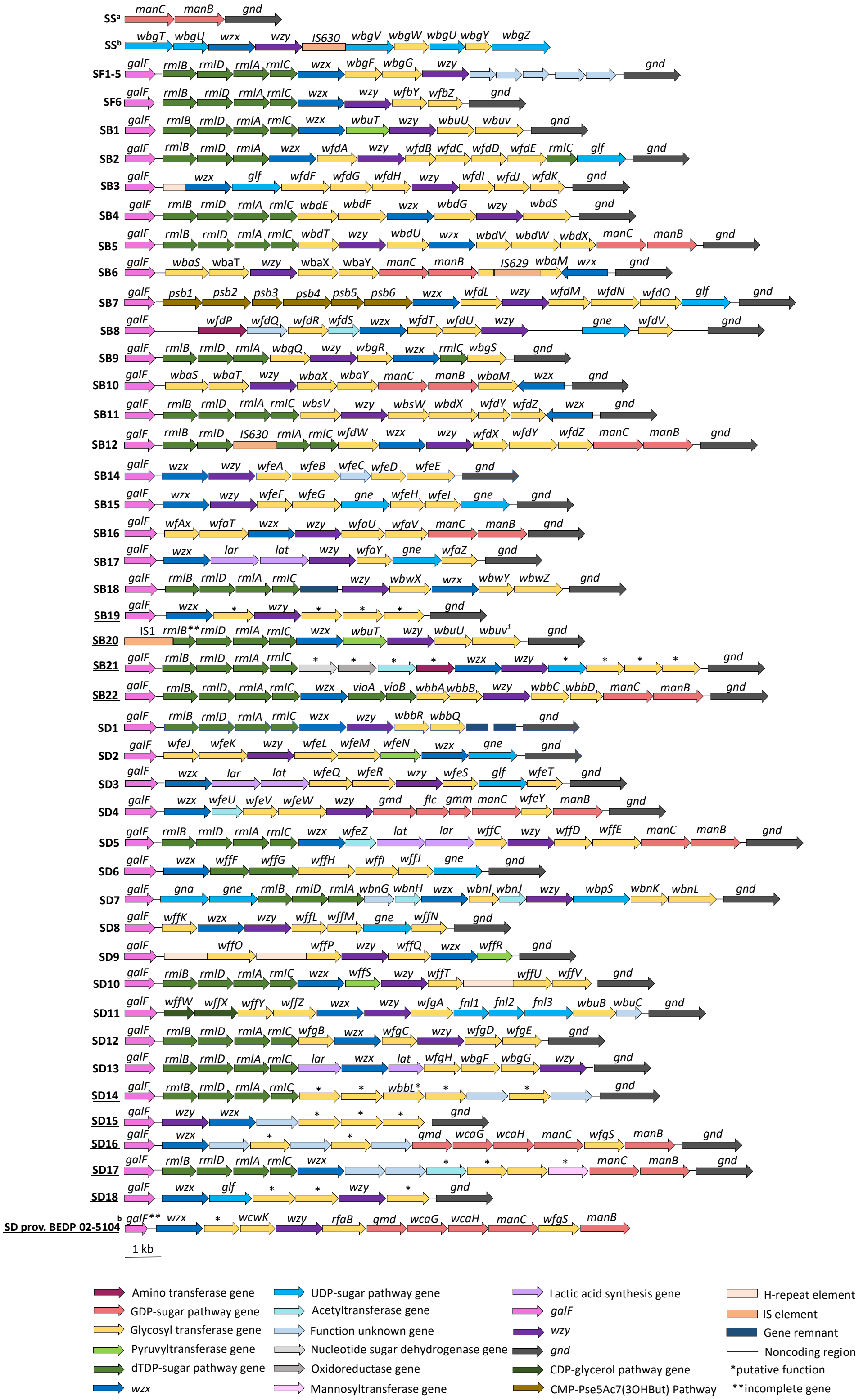

**Supplementary Figure 6:** Representation of all previously and newly characterised *Shigella* O-antigen gene clusters. The names underlined correspond to those presented for the first time. Open arrows represent the location and orientation of putative genes. SS[a] is a chromosomal remnant *rfb* from *S. sonnei*. A b in superscript indicates that the cluster is located on a plasmid. A 1 in superscript indicates that the gene may be truncated by an IS*1* in some strains.

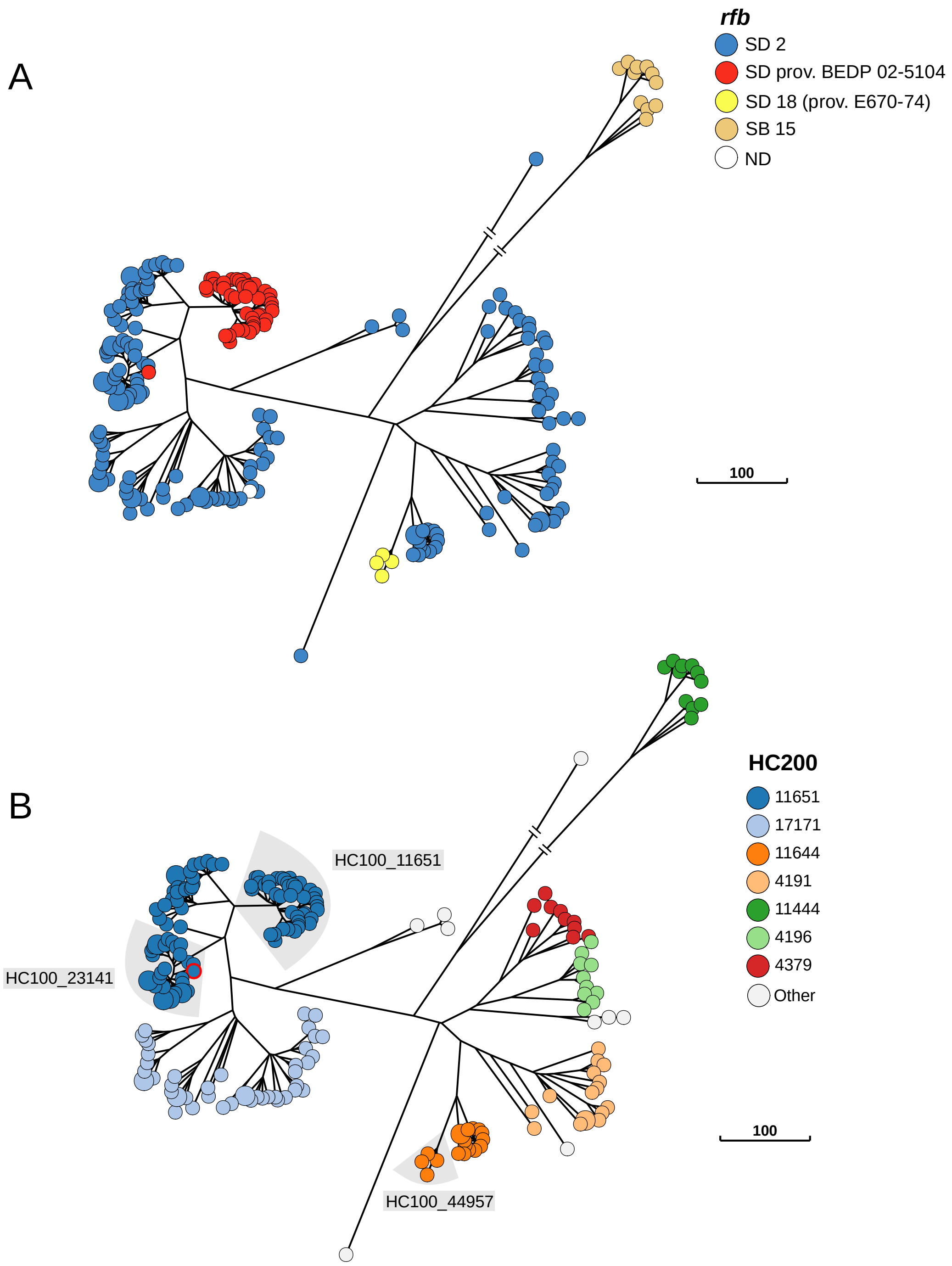

**Supplementary Figure 7:** NINJA neighbour-joining GrapeTree based on the cgMLST data for all S2d genomes (HC1100_4191, *n* = 222) present in EnteroBase (May 06 2021). (A) The tree nodes are colour-coded by type of *rfb* gene. ND indicates that the *rfb* cluster could not be identified. (B) The tree nodes are colour-coded by HC200 data. HC200 groups with fewer than three isolates are represented by white nodes. All *S. dysenteriae* prov. BEDP 02-5104 fall within HC100_11651, except for one strain (highlighted in red).
